# Chromatin interaction maps of human arterioles reveal new mechanisms for the genetic regulation of blood pressure

**DOI:** 10.1101/2024.10.09.617511

**Authors:** Yong Liu, Rajan Pandey, Qiongzi Qiu, Pengyuan Liu, Hong Xue, Jingli Wang, Bhavika Therani, Rong Ying, Kristie Usa, Michael Grzybowski, Chun Yang, Manoj K. Mishra, Andrew S. Greene, Allen W. Cowley, Sridhar Rao, Aron M. Geurts, Michael E. Widlansky, Mingyu Liang

## Abstract

Arterioles are small blood vessels located just upstream of capillaries in nearly all tissues. The constriction and dilation of arterioles regulate tissue perfusion and are primary determinants of systemic blood pressure (BP). Abnormalities in arterioles are central to the development of major diseases such as hypertension, stroke, and microvascular complications of diabetes. Despite the broad and essential role of arterioles in physiology and disease, current knowledge of the functional genomics of arterioles is largely absent, partly because it is challenging to obtain and analyze human arteriole samples. Here, we report extensive maps of chromatin interactions, single-cell expression, and other molecular features in human arterioles and uncover new mechanisms linking human genetic variants to gene expression in vascular cells and the development of hypertension. Compared to large arteries, arterioles exhibited a higher proportion of pericytes which were strongly associated with BP traits. BP-associated single nucleotide polymorphisms (SNPs) were enriched in chromatin interaction regions in arterioles, particularly through enhancer SNP-promoter interactions, which were further linked to gene expression specificity across tissue components and cell types. Using genomic editing in animal models and human induced pluripotent stem cells, we discovered novel mechanisms linking BP-associated noncoding SNP rs1882961 to gene expression through long-range chromatin contacts and revealed remarkable effects of a 4-bp noncoding genomic segment on hypertension in vivo. We anticipate that our rich data and findings will advance the study of the numerous diseases involving arterioles. Moreover, our approach of integrating chromatin interaction mapping in trait-relevant tissues with SNP analysis and in vivo and in vitro genome editing can be applied broadly to bridge the critical gap between genetic discoveries and physiological understanding.

## INTRODUCTION

Arterioles are small blood vessels found just upstream of capillary beds in nearly all tissues. Arterioles have diameters of 10 to 300 µm and consist of one layer of endothelial cells and one to a few layers of smooth muscle cells. The constriction and dilation of arterioles controls the blood flow to tissue regions. Therefore, arteriolar function is essential for maintaining proper distribution of blood across tissue regions at rest and changing blood flow in response to stimuli such as stress, exercise, and food ingestion. Additionally, arterioles are critical for determining systemic blood pressure (BP). BP is the product of total peripheral vascular resistance and cardiac output. Arterioles account for 60% to 70% of total peripheral vascular resistance in humans. Abnormalities in arterioles contribute to the development of major diseases such as hypertension, stroke, and many other microvascular complications of diabetes and hypertension^1–3^.

Despite the broad and essential role of arterioles in physiology and disease, current knowledge of the functional genomic features of arterioles is limited. This is in part because it is difficult to obtain arteriole samples for analysis, which requires skilled, microscopic tissue dissection^2^. Some functional genomic data is available for larger arteries with diameters of several mm or more. For example, the Encyclopedia of DNA Elements (ENCODE) project and the Genotype-Tissue Expression (GTEx) project include aorta, coronary artery, and tibial artery in their analysis^4,5^. However, larger arteries and arterioles are physiologically highly distinct. Larger arteries do not contribute to the regulation of tissue perfusion or BP as arterioles do, and their cellular composition and properties are likely different than arterioles^6,7^. Therefore, functional genomic insights obtained from the analysis of larger arteries may not be applicable to arterioles, leaving a critical knowledge gap.

In this study, we utilized our experience in arteriolar and functional genomic research^8–10^ to establish an extensive epigenomic landscape for human arterioles. We generated data for global and promoter-focused chromatin conformation, DNA methylation, and bulk and single-nucleus gene expression in human arterioles or tissue components of human arterioles. Integrated analysis of these datasets revealed novel insights into the molecular basis of arteriolar function and the genetic mechanisms underlying the regulation of arteriole-related traits including BP. We further investigated the BP-associated, noncoding single nucleotide polymorphism (SNP) rs1882961 that we found to form long-range chromatin interactions with *NRIP1* promoter. Using genome editing in animal models and human induced pluripotent stem cells (hiPSCs), we uncovered the mechanism by which rs1882961 regulated NRIP1 expression and demonstrated robust effects for a rs1882961 orthologous site on BP in a rat model.

## RESULTS

### The cellular composition and properties of human arterioles are distinct from larger arteries

We started by examining the cell type composition of human arterioles using single-nucleus RNA-seq (snRNA-seq). The arterioles were dissected from adipose tissues from surgical discards. Three arterioles, one from each subject, were pooled to provide sufficient material for snRNA-seq (**Supplementary Table S1**). The arterioles were 100-150μm in diameter and a few mm long.

Despite the small amount of input tissue, we obtained snRNA-seq data meeting our quality criteria from 1,205 nuclei. We detected several major cell types previously detected in the thoracic aorta using the same set of marker genes^11^ (**Fig. 1A, 1B**). The proportions of the cell types were substantially different between the arteriole and the aorta. To account for variable recovery rates for different cell types during nucleus isolation, we calculated proportions of cell types relative to endothelial cells in each tissue. The proportion of pericytes was higher, and the proportion of vascular smooth muscle cells lower, in the arteriole than the aorta (**Fig. 1C**). For each cell type, the Spearman correlation coefficients of gene expression profiles between the arteriole and the aorta ranged from 0.52 to 0.76 (**Fig. 1D**). We examined the enrichment of genes reported by genome-wide association studies (GWAS) of arteriole-related traits in each cell type. GWAS for diastolic BP, systolic BP, pulse pressure, hypertension, stroke, diabetic nephropathy, diabetic retinopathy, and peripheral arterial disease were included in this analysis. We found that BP-associated GWAS genes were enriched in several arteriolar and aorta cell types but to substantially different extents (**Fig. 1E**). Genes associated with diastolic BP were enriched in arteriolar, but not aortic, endothelial cells, and they were substantially more enriched in arteriolar pericytes than aortic pericytes. Genes associated with systolic BP and pulse pressure were enriched in aortic, but not arteriolar, smooth muscle cells and in arteriolar fibroblasts much more than aortic fibroblasts (**Fig. 1E**).

**Figure 1.**
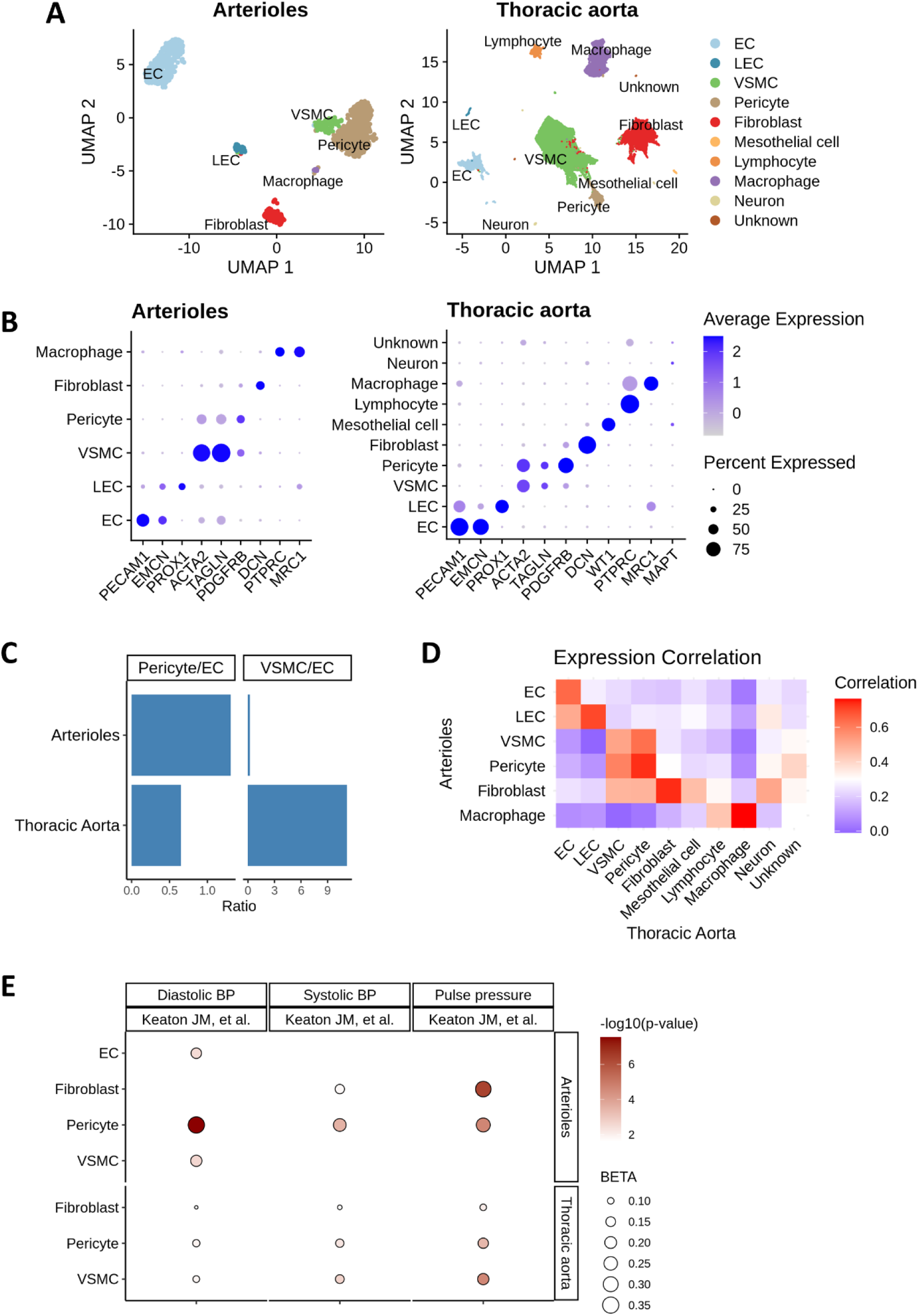
The cellular composition and properties of human arterioles exhibit distinct features compared with the aorta. **A.** UMAP plot showing major cell types identified in arteriole and thoracic aorta. **B.** Dot plot displaying marker gene expression across different cell types. **C.** Relative proportions of pericytes to ECs and VSMCs to ECs in arteriole and thoracic aorta, shown separately. **D.** Heatmap showing pairwise gene expression correlations across cell types in arteriole and thoracic aorta. **E.** Dot plot showing significant associations between cell types and blood pressure-related traits from MAGMA cell typing analysis. Cell type abbreviations: EC (endothelial cell), LEC (lymphatic endothelial cell), VSMC (vascular smooth muscle cell).

These findings indicate that the cellular composition and properties of human arterioles differ substantially from large arteries, suggesting that many of the epigenomic insights gained from analyzing large arteries may not apply to arterioles.

### Chromatin interactions, transcriptome, and DNA methylation in human arterioles

We constructed reference profiles of chromatin interactions, gene expression, and DNA methylation in human arterioles. The source subjects included a mix of race and sex (**Supplementary Table S1**). From each subject, one or more arterioles were separated into endothelial cells and endothelium-denuded arterioles (EDA), and one or more were analyzed as intact arterioles without the separation. The chromatin interaction analysis required more input material than what was available from a single subject. Therefore, arteriolar tissues from several subjects were combined to generate a reference chromatin interaction map for each tissue component.

Two chromatin interaction assays were performed: a Micro-C assay for global mapping and a pan-promoter capture Micro-C assay for mapping chromatin interactions with promoters of more than 27,000 coding and noncoding genes.

The Micro-C assay detected approximately 1,000 chromatin loops at 4-kbp resolution and close to 5,000 loops at 8-kbp or 16-kbp resolution in intact arterioles and EDAs from 1.4 billion and 1.7 billion read pairs, respectively (**Supplementary Tables S2-S9**). All loops were intrachromosomal by the default settings of the analysis. The median loop size ranges from 96 kbp at 4-kbp resolution to nearly 500 kbp at 16-kbp resolution (**Supplementary Fig. S1**). Substantial overlaps, defined by overlap of both contact regions that defined a loop, were observed between loops detected at different resolutions for a given tissue type and between loops detected in the two tissue types, supporting the robustness of the loop identification (**Fig. 2A**). For example, at the 16 kbp resolution, 33% of all detected loops were shared between intact arterioles and EDAs (**Fig. 2A**). Only a few dozen loops were detected in arteriolar endothelial cells, suggesting that, even with pooling, the amount of material in the arteriolar endothelial cell sample was insufficient for detecting most chromatin loops.

**Figure 2.**
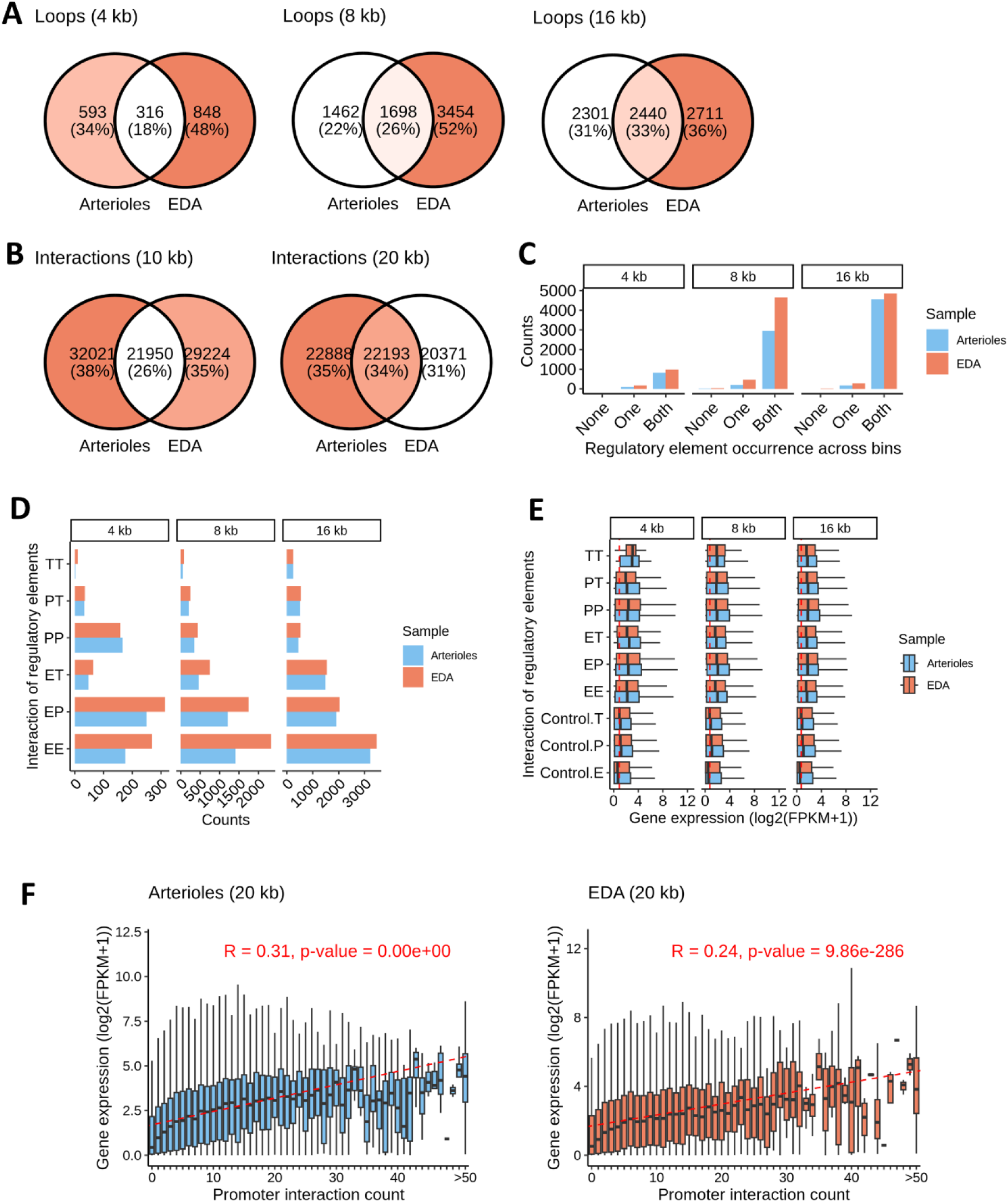
Chromatin contact regions in human arterioles contain DNA regulatory elements and correlate with greater gene expression. **A.** Venn diagrams showing the overlap between loops identified in arterioles and EC-denuded arterioles (EDA) by Micro-C at 4 kb, 8 kb, and 16kb resolutions. **B.** Venn diagrams showing the overlap between interactions identified in arterioles and EDA by pan-promoter Capture Micro-C at 10 kb and 20 kb resolutions. **C.** Bar plot showing the number of loops with regulatory elements present in one or both chromatin contact regions in arterioles and EDA across resolutions. “None,” “one,” and “both” indicate the presence of regulatory elements in neither, one, or both interacting regions, respectively. **D.** Number of chromatin interactions classified by interaction type: promoter-promoter (PP), enhancer-promoter (EP), enhancer-enhancer (EE), enhancer-transcription factor binding site (ET), promoter-transcription factor binding site (PT), and transcription factor binding site-transcription factor binding site (TT). Categories are not mutually exclusive. **E.** Expression levels of genes proximal to chromatin interactions (within 5 kb upstream or downstream of contact regions), grouped by interaction type (PP, EP, EE, ET, PT, TT). Controls include genes proximal to regulatory regions, including promoters, enhancers, or transcription factor binding sites, that were not in contact regions based on our data. **F.** Boxplot showing gene expression levels grouped by the number of chromatin interactions involving gene promoter regions. Spearman correlation was applied.

The pan-promoter capture Micro-C assay detected more than 100,000 and 80,000 chromatin interactions with promoters at 10-kbp and 20-kbp resolution, respectively, in intact arterioles and EDAs (**Supplementary Table S2, S10-S14**). Approximately 70% of these contacts were intrachromosomal. Approximately 50,000 intrachromosomal promoter interactions in each tissue with distances between chromatin contacts from 10 kbp to 2 million bp were included in the downstream analysis. Chromatin interactions were detected for the promoters of approximately 70% of the 20,089 protein-coding genes covered by the human pan-promoter probe panel. Chromatin interactions were also detected for the promoters of approximately 30% of the 43,051 nonprotein-coding genes defined by Ensembl GRCh38 reference dataset. The median distance of the interacting regions was approximately 100 kbp (**Supplementary Fig. S2**). Substantial overlaps, defined by overlap of both contact regions, were observed between interactions detected in the two tissue types. At 10- and 20-kbp resolution, 26% to 34% of the interactions were shared between intact arterioles and EDAs (**Fig. 2B**). More interactions were detected at 1-, 2-, or 5-kbp resolution, but the percent of intrachromosomal interactions were below 35% and the overlaps between the two tissues were less than 2%, suggesting greater noise in the result at these resolutions. No significant interactions were detected in the arteriolar endothelia sample likely because of, again, insufficient input material.

Poly(A)-dependent bulk RNA-seq analysis detected approximately 30,000 genes in intact arterioles, EDAs, and arteriolar endothelial cells (n=3 each) (**Supplementary Table S2, Fig. S3**). As expected, the transcriptome profiles of arteriolar endothelial cells differed substantially from the intact arterioles and EDAs, with 2,831 and 2,818 differentially expressed genes, respectively (**Supplementary Fig. S3**). The arteriolar endothelial cells were enriched for endothelial marker genes, while the arterioles and EDAs were enriched for marker genes for vascular smooth muscles (**Supplementary Fig. S4**). 226 genes were differentially expressed between arterioles and EDAs.

Several hundred differentially methylated regions were identified between arteriolar endothelial cells and intact arterioles or EDAs, and a few dozen identified between intact arterioles and EDAs (**Supplementary Table S2**; **Supplementary Fig. S5**). More than half of these differentially methylated regions were in gene promoter regions (**Supplementary Fig. S5**). Similar to previous reports^12,13^, the overall correlation between RNA abundance and gene promoter methylation was limited (**Supplementary Fig. S6**).

### Chromatin contact regions are enriched for regulatory elements and associated with higher gene expression

We obtained DNA regulatory elements including enhancers, promoters, and transcriptional factor binding sites from Ensembl^14^. More than 90% of the chromatin loops in arterioles and EDAs, defined by global Micro-C, contained regulatory elements in both contact regions, close to 8% contained regulatory elements in one of the two contact regions, and only a small fraction did not contain any regulatory elements in their contact regions (**Fig. 2C**). The overlap of chromatin contact regions and regulatory elements was substantial as only 12% to 18% of the genome was covered by the contact regions and 16% covered by the regulatory elements.

We grouped the chromatin loops in arterioles and EDAs into those involving enhancer-enhancer (EE), promoter-promoter (PP), enhancer-promoter (EP), enhancer-transcription factor binding site (ET), promoter-transcription factor binding site (PT), and transcription factor binding site-transcription factor binding site (TT) interactions (**Fig. 2D)**. All types of chromatin contacts were associated with higher expression of nearby genes (**Fig. 2E**).

Based on the pan-promoter capture Micro-C analysis, the vast majority (89-96%) of chromatin regions interacting with gene promoters contained known regulatory elements (**Supplementary Fig. S7A**). Approximately 72% of the chromatin regions interacting with gene promoters contained enhancers, and 15% contained transcriptional factor binding sites (**Supplementary Fig. S7B**). RNA abundance was highly significantly correlated with the number of chromatin regions interacting with gene promoters (**Fig. 2F**). The average number of chromatin interactions was 3.8 per gene.

### Representative genes essential for arteriolar function

We examined promoter chromatin interactions, DNA methylation, and RNA abundance for several genes essential for arteriolar function. *AGTR1* (Angiotensin II Receptor Type 1) had greater expression in intact arterioles and EDA than in arteriolar endothelial cells (**Fig. 3A**). We detected interactions of *AGTR1* promoter with intergenic genomic segments approximately 100 kbp and 50 kbp upstream in intact arterioles and EDA, respectively (**Fig. 3A**). One of these interacting regions contained known enhancers based on Ensembl annotation. In EDA, *AGTR1* promoter also interacted with a genomic region nearly 600 kbp downstream of *AGTR1* (**Fig. 3A**). This genomic region contained known enhancers and binding sites for transcriptional factors and the CCCTC-binding factor (CTCF) important for chromatin conformation and transcriptional regulation.

**Figure 3.**
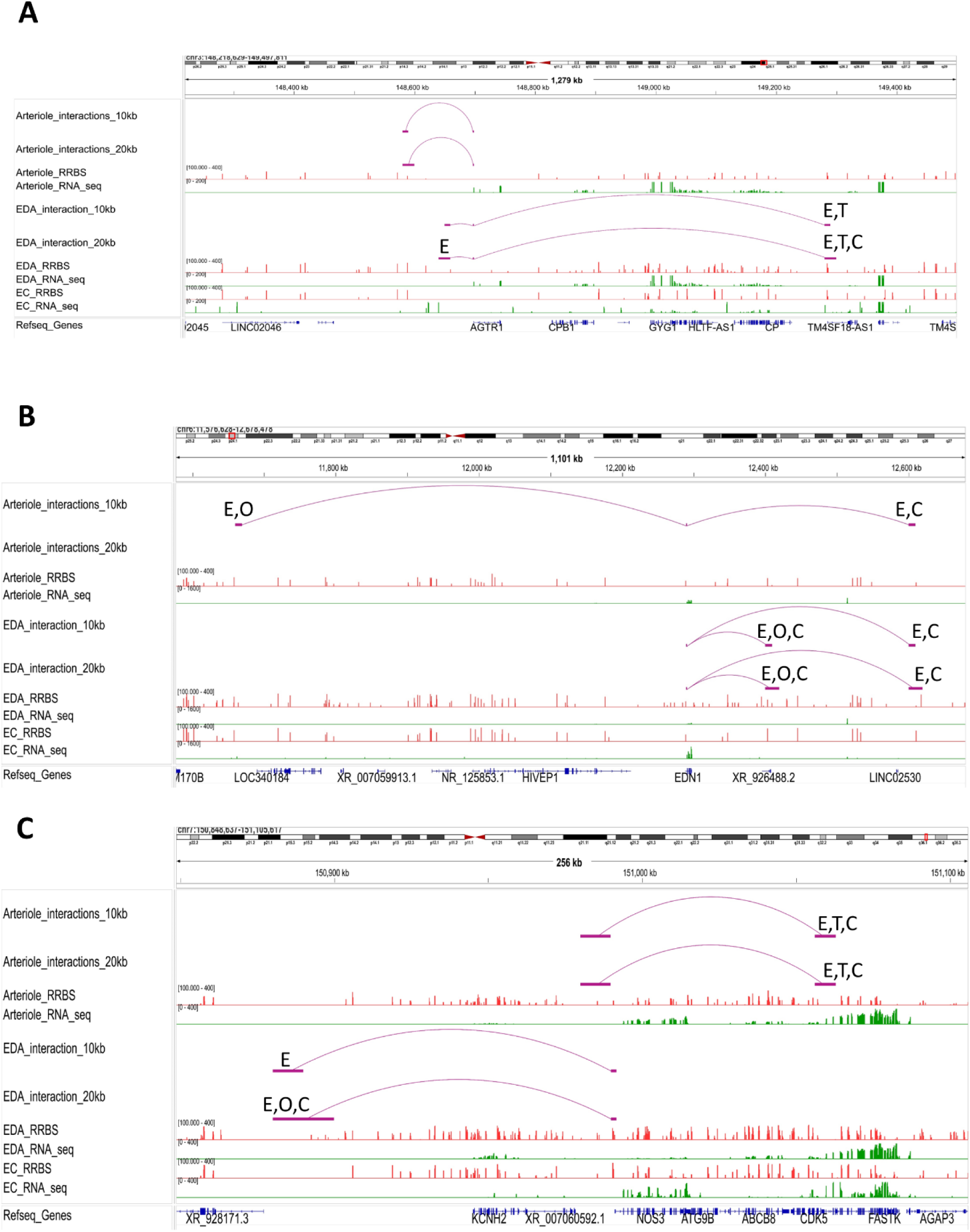
Chromatin interactions with gene promoters, DNA methylation, and mRNA abundance of representative genes essential to arteriolar function. **A.** *AGTR1* (Angiotensin II Receptor Type 1). **B.** *EDN1* (Endothelin 1). C. *NOS3* (Nitric Oxide Synthase 3). Linux-based IGV was used to plot chromatin interactions based on pan-promoter capture Micro-C, RNA abundance based on poly(A)-dependent RNA-seq, and methylation levels based on RRBS. Only chromatin interactions with the promoter of the gene of interest are shown. The chromatin contact regions were compared with the regulatory elements as defined by Ensembl for overlaps. E, enhancer; T, transcriptional factor binding site; C, CCCTC-binding factor (CTCF) binding site; O, open chromatin region.

In contrast, *EDN1* (Endothelin-1) and *NOS3* (Nitric Oxide Synthase 3) transcripts were abundant in intact arterioles and endothelial cells but much less so in EDA (**Fig. 3B, 3C**). For *EDN1* promoter, interactions with a genomic segment approximately 300 kbp downstream of the gene were detected in both arterioles and EDA (**Fig. 3B**). Additional interacting regions approximately 600 kbp upstream and 100 kbp downstream of the gene were detected in arterioles and EDA, respectively. All these interacting regions contained known enhancers, open chromatin regions, or binding sites for CTCF. The genomic region surrounding *EDN1* had more DNA methylation in EDA than in arterioles and endothelial cells (**Fig. 3B**). For *NOS3* promoter, interacting regions were detected approximately 150 kbp downstream and 200 kbp upstream in arterioles and EDA, respectively, and these regions contained known enhancers, binding sites for transcriptional factors and CTCF, or open chromatin regions (**Fig. 3C**). The *NOS3* gene body had more DNA methylation in EDA than in arterioles and endothelial cells.

These characteristics indicate that chromatin interactions and DNA methylation may play a here-to-fore unrecognized role in regulating the expression of genes essential to arteriolar function, suggesting new future directions for the study of these genes.

### Chromatin contact regions in human arterioles are enriched for BP-associated SNPs

To investigate the relation between arteriolar chromatin architecture and genetic determinants of arteriole-related phenotypes, we integrated the Micro-C and pan-promoter capture Micro-C data with an analysis of GWAS SNPs associated with arteriole-related traits, including systolic BP, diastolic BP, pulse pressure, essential hypertension, stroke, peripheral arterial disease, diabetic nephropathy, and diabetic retinopathy. By examining the overlap between these SNPs and chromatin interaction regions in both intact arterioles and EDA, we observed that SNPs associated with systolic BP, diastolic BP and pulse pressure were consistently enriched in chromatin contact regions compared to SNPs linked to non-arteriole-related traits (**Fig. 4A**; Methods). Additionally, stroke-associated SNPs were significantly enriched in chromatin contacts in intact arteriole but not in EDA, consistent with a prominent role of endothelial cells in stroke pathogenesis. Our arteriole-based Micro-C data showed a higher capture of BP-associated SNPs per 1Mbp loop contact region compared to Hi-C data from tibial arteries and aorta in ENCODE^4^ (**Fig. 4B**).

**Figure 4.**
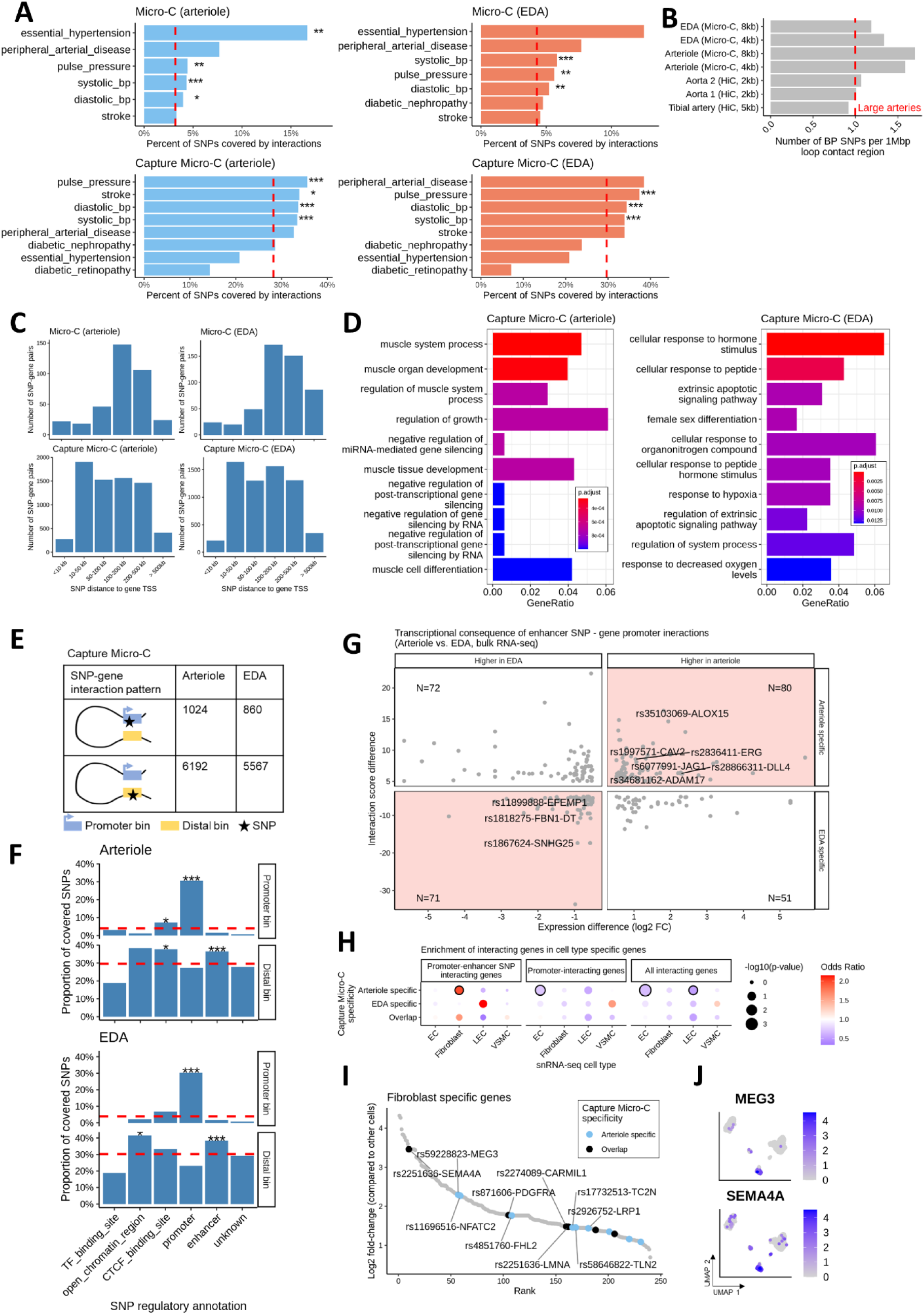
Chromatin contact regions in human arterioles are enriched for SNPs associated with blood pressure and stroke. **A.** Bar plot showing the percentage of SNPs covered by Micro-C or pan-promoter Capture Micro-C in arterioles and endothelium-denuded arteriole (EDA), grouped by different arteriole-related traits. Red dashed lines indicate the average SNP coverage rate from non-arteriole-related traits (Methods). A one-sided binomial test was used to assess the significance of the higher SNP coverage rates in arteriole-related traits. **B.** Bar plot showing the density of BP-associated SNPs within 1 Mb loop contact regions across our Micro-C data in arterioles and EDA, compared to Hi-C data from the tibial artery and aorta (ENCODE). Red dashed lines represent the average SNP density from the Hi-C data of the tibial artery and aorta. **C.** Bar plot showing the distance distribution of SNP-gene pairs capture by Micro-C or pan-promoter Capture Micro-C in arteriole and EDA, grouped by distance categories. SNPs associated with arteriole-related traits (as listed in Fig. 4A) were included. **D.** Pathway enrichment results of genes associated with arteriole-related SNPs through chromatin interactions captured by pan-promoter Capture Micro-C, shown separately for arteriole and EDA. **E.** Schematic of SNP-gene interaction patterns grouped by SNP bin location in pan-promoter Capture Micro-C, along with the corresponding interaction numbers in arterioles and EDA. **F.** Percentage of covered SNPs based on the regulatory elements in which the SNPs are located, calculated separately for promoter bins and distal bins. Red dashed lines represent the overall coverage rate of all SNPs. A Fisher’s exact test was used to assess the significance of SNP coverage in specific regulatory groups, followed by Benjamini-Hochberg (BH) correction for p-values. Only significant results with an odds ratio > 1 are highlighted with asterisks. **G.** Dot plot showing the relationship between enhancer SNP-gene promoter interactions and corresponding gene expression. Only interactions specifically identified in arterioles or EDA, and genes with significant expression differences between arterioles and EDA (BH-corrected p-value < 0.05, absolute log2 fold change > 0.5) were included. Consistent results are highlighted with a red background. Notable SNP-gene pairs are labeled, and the numbers for SNP-gene pairs in each condition are shown in the corners of each quadrant. **H.** Dot plot showing the enrichment of arteriole-related SNP-interacting genes in cell type-specific genes using a Fisher test. SNP-interacting genes were grouped by interaction type, including promoter-enhancer SNP-interacting genes, promoter-interacting genes, and all-interacting genes (faceted by columns), and by the occurrence of interactions in arterioles only, EDA only, or both (faceted by rows). **I.** Rank plot of genes significantly higher expressed in fibroblast compared to all other cells (Bonferroni-corrected p-value < 0.05 by Wilcoxon test, ordered by log2 fold change). Genes having chromatin interactions involving arteriole-related SNPs, identified via pan-promoter Capture Micro-C, were highlighted and colored based on the occurrence in samples. Top 10 genes were label. **J.** UMAP plots showing the expression pattern of MEG3 and SEMA4A across cell types in arterioles, based on snRNA-seq data. Statistical significance is indicated as follows: *p < 0.05, **p < 0.01, and **p < 0.001.

Further analysis of SNPs associated with arteriole-related traits and genes interacting with these SNPs captured by our chromatin contacts identified an average of 344 long-distance SNP-gene interactions (>100 kb) in the Micro-C data and an average of 3,331 interactions in the pan-promoter capture Micro-C data (**Fig. 4C**). Functional enrichment analysis of genes interacting with SNPs via chromatin contacts, based on the pan-promoter capture Micro-C data, showed significant enrichment in muscle system processes and tissue development in arterioles. In the EDA dataset, enriched pathways were related to cellular response to external stimuli, including hormone and peptide responses, apoptotic signaling and hypoxia (**Fig. 4D**).

### Enhancer SNP-promoter interactions linked to gene expression variation in arterioles and EDA

In our pan-promoter capture Micro-C data, each interaction includes a captured promoter bin and a distal interacting bin (**Supplementary Table S11-S14**). Approximately 87% of the SNPs in the captured SNP-gene pairs were in the distal interacting bin (6192 out of 7216 and 860 out of 6472) (**Fig. 4E**). These SNPs were significantly enriched in enhancer regions, suggesting that SNPs frequently exert their influence on gene promoter via enhancer regions when mediated by chromatin interactions (**Fig. 4F**). To assess potential impact of enhancer SNPs interacting with gene promoters on downstream gene expression, we selected SNP-gene pairs where these interactions were uniquely detected in either intact arterioles or EDAs. We focused our analysis on genes with significant expression differences (absolute log2 fold change > 0.5) to specifically assess the role of chromatin interactions on notable gene expression variations. We identified 151 SNP-gene pairs where the specific SNP-gene interaction and significantly higher gene expression were both observed in the same tissue (**Fig. 4G**). This analysis revealed notable SNP-gene pairs. In intact arteriole but not EDA, several blood pressure-related SNPs were linked to endothelial cell (EC)-associated genes, including rs2836411-*ERG* (previously mapped in GWAS Catalog and confirmed by our data), rs28866311-*DLL4*, and rs34681162-*ADAM17* (both newly identified). *DLL4* is involved in angiogenesis via Notch signaling^15^, while EC-derived *ADAM17* has been linked to dysregulated coronary arteriole dilation^16^. *ERG* regulates angiogenesis and vascular stability by promoting VE-cadherin expression and Wnt/β-catenin signaling, ensuring endothelial integrity and survival^17,18^. EDA-specific interactions involved *FBN1-DT*, a divergent transcript of *FBN1*. Mutations in *FBN1* are linked to Marfan syndrome, which involves vascular abnormalities, including VSMCs-related issues^19^.

### Arteriole enhancer SNP-promoter interactions enriched in fibroblast-specific genes

To investigate whether SNP-gene interactions are associated with cell type-specific gene expression patterns, we performed enrichment analysis of arteriole-specific, EDA-specific, and overlapping SNP-interacting genes against cell type-specific gene expression profiles from arteriole snRNA-seq data. We found that arteriole-specific promoter-enhancer SNP interacting genes were significantly enriched in genes highly expressed in fibroblasts, a unique pattern compared to other interaction modes, including promoter-interacting genes and all interacting genes (**Fig. 4H**). By mapping SNP-interacting genes to fibroblast-specific gene expression, we identified several top enhancer SNP-gene promoter interactions, including rs59228823-*MEG3* and rs2251636-*SEMA4A* (**Fig. 4I**). Both genes showed fibroblast-specific expression, suggesting a potential cell type-specific regulatory effect of the linked SNPs (**Fig. 4J**). Collectively, these findings showed the strength of our dataset in uncovering how chromatin interactions bridge the genetic susceptibility to arterial-related phenotypes with underlying cellular mechanisms.

### BP-associated SNP rs1882961 interacts with *NRIP1* promoter in human arterioles and regulates NRIP1 expression

The above findings suggested that SNPs might influence arteriole-related phenotypes by regulating genes whose promoters interacted with the SNPs across long genomic distances. In a proof of principle study, we experimentally investigated a selected BP-associated SNP. There were 8,790 pairs of arteriolar trait-associated SNPs and genes whose promoters interacted with the genomic regions containing these SNPs based on our arteriolar and EDA pan-promoter capture Micro-C dataset. We filtered these pairs based on both biological and technical criteria (**Fig. 5A**). The biological criteria included BP-associated SNPs, promoters of protein-coding genes, low promoter methylation levels, and RNA abundance above a minimum threshold. Technical criteria include not being in linkage disequilibrium with other SNPs for ease of genomic editing, cross-species conservation of the SNP genomic region to facilitate studies in animal models, and genes paired with only one SNP to avoid confounding effects from multiple variants. Furthermore, we focused on SNP-gene pairs for which the SNP is more than 100 kbp away from the gene as it is particularly challenging to identify the effector genes for these SNPs and understand the genomic mechanisms involved. These filters narrowed the list of SNP-gene pairs down to 89, from which we performed experimental studies on rs1882961 and *NRIP1*.

**Figure 5.**
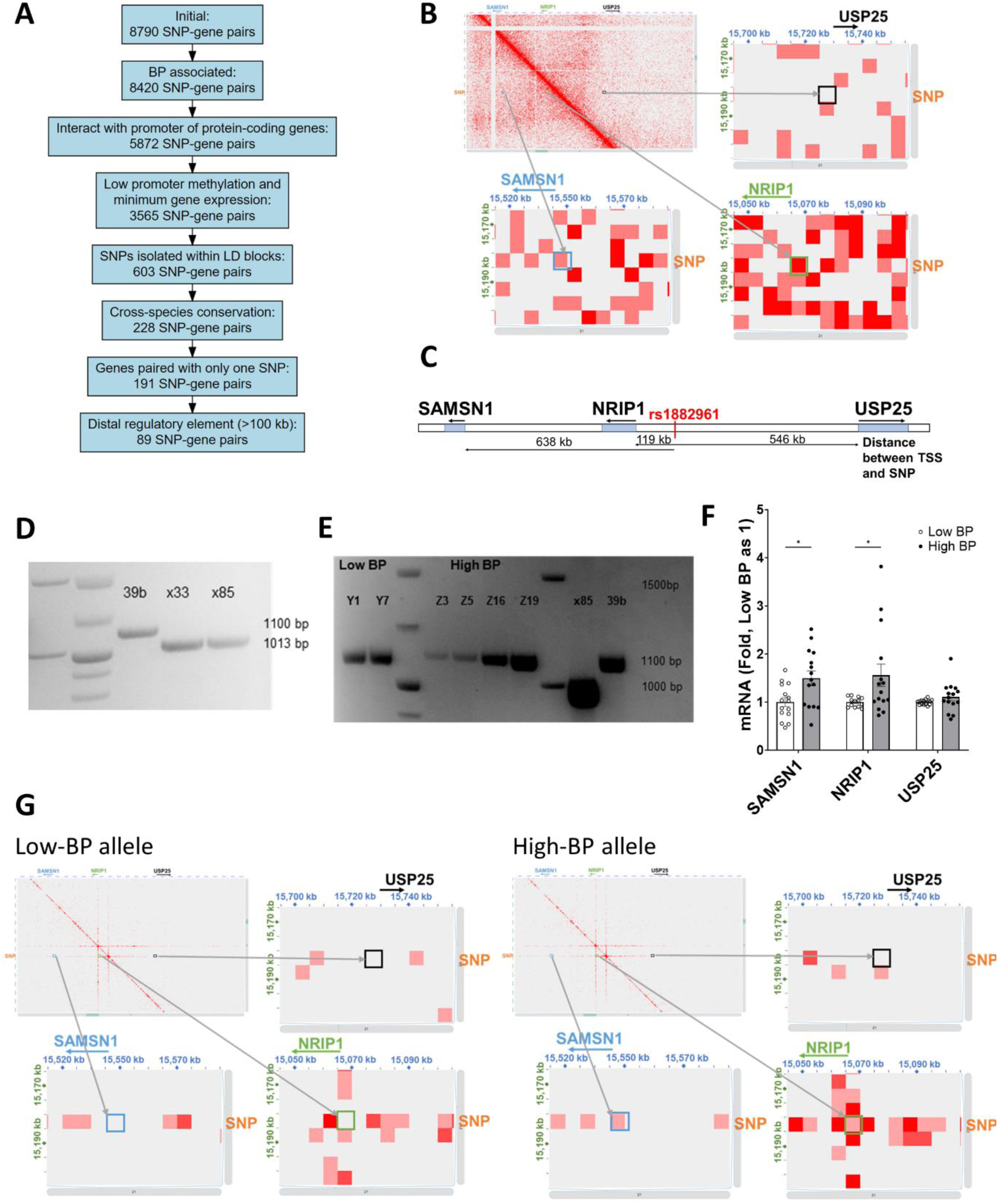
BP-associated noncoding SNP rs1882961 interacts with NRIP1 promoter 119 kbp away and influences NRIP1 expression. **A.** Prioritization of SNP-gene pairs for experimental analysis. **B.** The genomic region containing rs1882961 interacted with NRIP1 promoter 119 kbp away from it in human endothelium-denuded arterioles, based on pan-promoter capture Micro-C analysis. **C.** Local gene organization around rs1882961 in the human genome. **D.** Deletion of DNA segment containing rs1882961. 39b, original iPSC cell line; X33 and X85, rs1882961-deleted cell lines. **E.** Reconstitution of the rs1882961 locus containing either homozygous low BP allele or high BP allele. 39b, original iPSC cell line; X85, rs1882961-deleted cell line; Y series, cell lines with reconstituted low-BP rs1882961 allele; Z series, cell lines with reconstituted high-BP rs1882961 allele. See Supplementary Fig. S8 for sequence confirmation. **F.** Homozygous high-BP allele of rs1882961, reconstituted in hiPSCs, increased NRIP1 and SAMSN1 expression in isogenic hiPSC-derived vascular smooth muscle cells (iVSMCs). N = 15; *, p < 0.05, unpaired t-test. **G.** Region-capture Micro-C analysis showed greater chromatin interactions between the rs1882961 region and the promoters of SAMSN1 and NRIP1 in iVSMCs with the high-BP allele of rs1882961, compared to iVSMCs with the low-BP allele. Each square color box in the zoom-in images is 5 kbp x 5 kbp.

rs1882961 is associated with systolic BP^20^. The genomic region containing rs1882961 interacted with *NRIP1* promoter region in EDA (**Fig. 5B**). The transcription start site of *NRIP1* is 119 kbp from rs1882961 (**Fig. 5C**). *NRIP1* encodes nuclear receptor interacting protein 1, also known as receptor-interacting protein 140 (RIP14). RIP140 is a coregulatory for most nuclear receptors including retinoic acid receptor, estrogen receptor, and thyroid hormone receptor, many of which could influence vascular function^21^. However, the involvement of *NRIP1* in blood pressure regulation and the role of rs1882961 in the regulation of *NRIP1* are unknown.

We experimentally examined the allelic effect of rs1882961 on the expression of *NRIP1* and other local genes *SAMSN1* and *USP25* in human vascular smooth muscle cells (**Fig. 5C**). To that end, we generated isogenic hiPSCs precisely edited to contain homozygous rs1882961-C (low-BP allele) or rs1882961-T (high-BP allele) (**Fig. 5D, 5E**; **Supplementary Fig. S8, S9**). The edited hiPSCs, three clones each, were differentiated to vascular smooth muscle cells (iVSMCs). In iVSMCs, NRIP1 and SAMSN1 were significantly upregulated in cells with the high-BP allele of rs1882961 compared with the low-BP allele (**Fig. 5F**). USP25 was not differentially expressed. We performed region-capture Micro-C analysis to capture any genomic segment interacting with rs1882961 or promoters of its neighboring genes. The analysis revealed greater chromatin interactions between the rs1882961 genomic region and the promoter regions of *NRIP1* and *SAMSN1*, but not *USP25*, in iVSMCs with the high-BP allele than the low-BP allele (**Fig. 5G**).

### A 4-bp segment encompassing the rs1882961 orthologous site alters BP in SS rats

We tested the effect of the rs1882961 genomic site on BP and Nrip1 expression in vivo. We mapped rs1882961 to position chr11:15,091,368 in the rat genome, which is 112 kbp from the TSS of Nrip1 (**Fig. 6A**). We successfully deleted a 4-bp segment that encompassed the rs1882961 orthologous site from the genome of the Dahl salt-sensitive (SS) rat using CRISPR/Cas9 (**Fig. 6A; Supplementary Fig. S10**). The mutant strain was designated SS-Δrs1882961. We phenotyped female rats as the hiPSCs used in the study above were derived from a female subject. Mean arterial pressure, systolic BP, diastolic BP, and pulse pressure, but not heart rate, in SS-Δrs1882961^−/−^ rats was significantly lower than wild-type littermate SS rats after the rats had been fed a high-salt (4% NaCl) diet (**Fig. 6B-5F**). The difference of mean arterial pressure occurred primarily in the dark phase (i.e., active phase) in a light-dark cycle, reaching approximately 15 mmHg during part of the dark phase (**Fig. 6G**). Nrip1 expression in mesenteric tissue was up-regulated in response to the high-salt diet but to a lesser extent in SS-Δrs1882961^−/−^ rats, resulting in significantly lower expression in SS-Δrs1882961^−/−^ rats than in wild-type SS rats on the high-salt diet (**Fig. 6H**). These studies demonstrate that a 4-bp segment of noncoding DNA encompassing the rs1882961 orthologous site regulates the expression of Nrip1 112 kbp away in vivo and influences the development of salt-induced hypertension.

**Figure 6.**
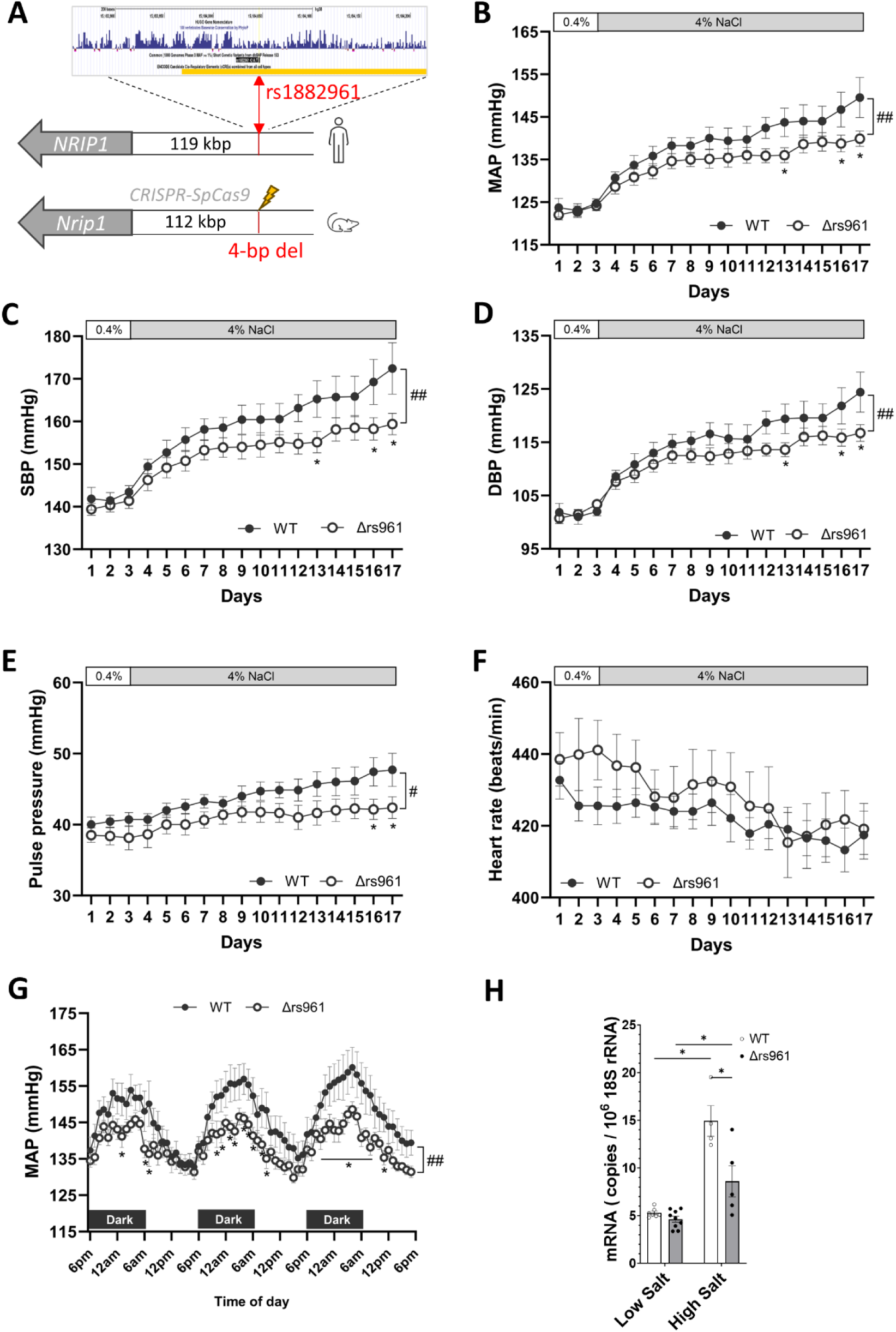
Deletion of a 4-bp noncoding genomic segment containing the rs1882961 orthologous site attenuates salt-induced hypertension in the Dahl salt-sensitive (SS) rat. **A.** Comparative mapping and deletion of a 4-bp rat genomic segment containing the rs1882961 orthologous site. See Supplementary Fig. S10 for additional detail and confirmation of deletion. The deletion of the 4-bp noncoding genomic segment in the SS rat attenuated salt-induced hypertension. **B.** Mean arterial pressure (MAP). **C.** Systolic blood pressure (SBP). **D.** Diastolic blood pressure (DBP). **E.** Pulse pressure. **F.** Heart rate. **G.** Hourly averages of MAP on days 12 to 14 on a 4% NaCl high salt diet. WT, wild-type littermate SS rats; Δrs961, SS-Δrs1882961#x2212;/− rats. N = 7 for WT and 8 for Δrs961. #, p < 0.05, ##, p < 0.01 for WT vs. Δrs961 by two-way RM ANOVA; *, p < 0.05 vs. WT by Holm-Sidak test. **H.** Nrip1 expression in the mesentery was lower in SS-Δrs1882961#x2212;/− rats compared to WT. N = 4-9. Data shown as Mean ± SEM. *, p<0.05, Two-way ANOVA followed by Holm-Sidak test.

## DISCUSSION

Long-range chromatin interactions play an important role in the regulation of gene expression^22,23^. We have found that, in human arterioles, regulatory elements are prevalent in chromatin contact regions and the frequency of long-range chromatin interaction with gene promoters is highly significantly correlated with RNA abundance. These findings indicate that long-range chromatin interaction may be a here-to-fore unrecognized mechanism underlying the essential physiological function and broad pathophysiological relevance of arterioles. We envision that our extensive dataset will drive forward mechanistic studies of hypertension, stroke, microvascular complications of diabetes and hypertension, and other arteriole-related diseases in new directions.

Long-range chromatin interactions are especially relevant to understanding the mechanisms by which noncoding sequence variants identified by GWAS influence their associated traits as many GWAS variants are located 10s of kbp from their nearest genes^24,25^. The approach that we used in this study integrates in-depth chromatin conformation analysis in trait-relevant human tissues with analysis of GWAS-nominated SNPs, in vivo and in vitro genome editing, and functional investigation. As demonstrated by our proof of principle study of rs1882961, this integrated approach is powerful and can be applied broadly to bridge the critical gap between genetic discoveries and physiological understanding.

It is remarkable that deletion of a mere 4 bp at a genomic site that is more than 100 kbp from the transcription start site of the nearest protein-coding gene changes hourly BP by up to 15 mmHg. The high-salt dietary challenge and possibly the genomic background of the SS rat may have contributed to the uncovering of this remarkable BP effect. By extrapolation, the SNP rs1882961 and other GWAS-nominated sequence variants may have greater effects on BP and other associated traits in groups of humans with permissive genomic backgrounds and exposed to certain stressors than the typically minute effect reported by GWAS. In addition, such robust phenotypic changes, along with the changes in Nrip1 expression in the mutant rat and edited cells not only reenforce the significance of long-range chromatin interaction in genetic and molecular regulation but also support its relevance to the regulation of physiological function.

This study is one of the first to apply Micro-C and capture Micro-C techniques to analyze human tissues^26,27^. The data was robust, considering the extremely limited input material compared with studies using cultured cells. We performed our chromatin interaction analyses, as well as DNA methylation analysis, in bulk tissues. While techniques for analyzing chromatin conformation and DNA methylation in single nuclei have been reported^28–30^, they require much more input materials than available from human arterioles, and data from these assays is sparser than from bulk tissue assays. Moreover, single-nucleus methods are not available for promoter capture Micro-C. Meanwhile, the robustness of our bulk tissue data is enhanced by the relative simplicity of cell type composition in arteriole compared to more complex organs. Additionally, we reduced the tissue complexity by analyzing separate tissue components in the current study.

We analyzed arterioles isolated from subcutaneous fat. Pathological changes in subcutaneous vessel structure and physiology are well-established to reflect the influence of cardiovascular risk factors and prevalent cardiovascular disease^31–35^. Physiological similarities and differences exist between arterioles in subcutaneous fat and those in visceral fat or internal organs^32,36,37^. It would be valuable to analyze arterioles in other tissue depots in future studies. Nevertheless, the reference molecular profiles generated in the current study will help to drive studies of arteriole-related traits and diseases in general as our study of rs1882961 has demonstrated. In addition, we chose to use subcutaneous fat because of the relative ease of obtaining such specimens from human participants, including via a minimally invasive biopsy in healthy individuals^9^. This means our data will be directly relevant to large-scale studies of human arterioles in which the ease of tissue procurement is particularly important as well as to studies involving healthy individuals not undergoing any surgery.

## METHODS

### Human arterioles

Human arterioles were obtained from discarded subcutaneous adipose tissue from surgery as described with modifications^9^. All protocols were approved by the Medical College of Wisconsin Institutional Review Board (IRB). Subject inclusion criteria were age 21-89 and without major cardiovascular diseases except for hypertension. Exclusion criteria were coronary heart disease, heart failure, diabetes, dyslipidemia, COVID19 infection within one-year, current tobacco use, and cancer chemotherapy drug exposure. Immediately upon receiving the discarded tissue, arterioles were isolated in cold (4°C) HEPES buffer by removing the surrounding fat and connective tissue. These intact isolated arterioles were snap-frozen in liquid nitrogen and stored at −80°C until use. For arterioles undergoing endothelial cell scraping, the isolated arterioles were cut open along their longitudinal direction with fine scissors and fixed lumen side-up with fine pins on a peri dish coated with silicone gel. After washing two times with PBS, the endothelial layer at the top of the lumen side was scrapped with a blade. After scraping, the blade was rinsed in an Eppendorf tube containing 700 µl of PBS; 800 µl of PBS was used to rinse the scrapped lumen side and collected into the same Eppendorf tube. The tube was spun at 300g at 4°C for 10 minutes. The pellet with 100 µl of supernatant was snap-frozen in liquid nitrogen and stored at −80°C until use. The vascular tissue that remained after the scraping was also collected as endothelium-denuded arteriole (EDA).

### Micro-C assay and data analysis

Micro-C assay^27^ was performed using Dovetail® Micro-C Kit following manufacturer’s User Guide for Tissue and Blood to create Micro-C libraries. Briefly, tissues were ground into fine powder with CryoGrinder^TM^. The ground tissue was crosslinked with disuccinimidyl glutarate and formaldehyde sequentially. The genomic DNA was digested *in situ* with MNase Enzyme mix to 20%-40% mononucleosomes. Genomic DNA was released from cells and put through a sequential process of binding to chromatin capture beads, end polishing, bridge ligation, intra-aggregate ligation, crosslink reversal, and DNA purification. Library was made from the purified DNA and subject to sequencing with Novaseq 6000 platform for approximately 800M 150PE reads.

Pair-end sequencing reads were cleaned with Trim Galore to remove adaptor and low-quality reads. We used JuicerBox, which was a one-click pipeline for processing HiC or Micro-C data, to generate hic file for direct visualization^38^. We followed Dovetail Micro-C Data Processing Guide to generate .mcool file and used it for loop calling with Mustache^39^.

### Pan-promoter capture Micro-C assay and data analysis

Pan-promoter capture Micro-C library was created using the Micro-C libraries generated as described above, Dovetail® human Pan Promoter Enrichment panel, and Dovetail Target Enrichment kit. The panel included 161,144 probes designed to cover 84,643 promoters for 19725 protein-coding genes and 7,630 long noncoding genes in human based on Ensembl GRCh38 version 86. The libraries were sequenced using the Illumina NovaSeq sequencers.

The data was analyzed following Dovetail Promoter Panel Data Processing Guide to generate mapped.PT.bam file. Direct chromatin interactions were then identified by CHiCAGO at different resolutions with the mapped.PT.bam file^40^. To ensure our dataset aligned with the most current promoter definitions, we updated the transcript promoter annotations using the latest Ensembl GRCh38 version 112. Following this update, we retained 99% of the original chromatin interactions that captured promoter regions (**Supplementary Table S10**). These retained interactions were used in the subsequent analysis. Genomic overlap analysis was performed using the GenomicRanges R package (v1.50.2). Regulatory element annotations were downloaded from Ensembl. To maintain consistency with the promoter definition used in Dovetail, promoter regions in the pan-promoter Capture Micro-C data were defined as 1000 bp upstream and 500 bp downstream of transcript TSSs.

### Poly(A)-dependent RNA-seq and data analysis

Poly(A)-dependent RNA-seq experiment and data analysis were performed as described^9^. An in-house pipeline was used to analyze RNA-seq data. Briefly, raw RNA-Seq sequence reads were pre-processed using Trim Galore (v0.6.2) (http://www.bioinformatics.babraham.ac.uk/projects/trim_galore/). Adapters and reads with low quality (base quality < 20) were removed prior to further analysis. The trimmed sequence reads longer than 20bp were mapped to the human genome (hg38) using hisat2 (v2.2.1)^41^. Transcript assembly and quantification were performed using StringTie2 (v2.2.1)^42^, based on the GENCODE reference annotation (v38). R package DESeq2 was used to identify differentially expressed genes^43^.

### Reduced representation bisulfite sequencing (RRBS) and data analysis

DNA methylation profiles at single-base resolution were analyzing using multiplexed RRBS as we described^12,13,44^. Trimmed sequences were mapped to the human reference genome (hg38) using Bismark (v0.16.1)^45^. Then, metilene (Version 0.23) with two-dimensional Kolmogorov–Smirnov test (KST) was applied to identify methylation regions (MRs) de novo^46^. Methylation rate of each region was calculated as the average methylation rate of each site across the region. Differentially methylated regions (DMR) were identified based on the following criteria: a minimum mean methylation difference between groups of 0.05, a minimum of five CpGs within the identified regions, and an FDR < 0.05 from the KST.

### Single-nucleus RNA-seq and data analysis

Human arterioles were broken up using a beads method, and nuclei were extracted using the ’Frankenstein’ protocol. Briefly, the arterioles were broken up in 0.5 ml Nuclei EZ Lysis buffer with Precellys Evolution Touch and tissue homogenizing ceramic beads at a setting of 6,500rpm for 20 seconds. Another 3.5 ml Nuclei EZ Lysis buffer was added to tissue homogenized buffer and incubated for 5 minutes on ice before being filtered through a 70 µm strainer mesh. The extracted nuclei were washed once with Nuclei Wash and Suspension Buffer. Single-nucleus RNA-seq libraries were prepared using Chromium Single Cell 3’ Reagent Kits V3 (10x Genomics, Pleasanton, CA), following the manufacturer’s user guide. snRNA-seq libraries were sequenced using the Illumina NovaSeq sequencers to generate approximately 30 GB or 100 million paired reads of data per library.

Raw fastq files were processed using CellRanger (v7.0.0) to generate feature matrices using 10x Genomics reference genome assembly hg38 (refdata-gex-GRCh38-2020-A)^47^. DoubletFinder R package (v2.0.3) was used to remove potential doublets, with the doublet rate set to max(0.3, #cells*8*1e-6)^48^. Cells with less than 200 genes or a mitochondrial percentage larger than 20% were excluded from the analysis.

For quality-controlled snRNA-seq data, the gene count matrix was log-transformed and scaled. The top 2000 most variable genes were identified using the FindVariableFeatures function from Seurat R package (v4.3.0.1)^49^, and then were used for principal component analysis (PCA). Clustering was performed using the top 20 principal components and the Louvain algorithm, with resolution adjusted from 0.5 to 2 in increments of 0.5. The optimal resolution was selected based on Clustree (v0.5.1) analysis^50^. To annotate identified clusters, differentially expressed genes (DEGs) between clusters were identified by Wilcoxon rank sum test using FindAllMarkers function (logfc.threshold = 0.25, min.pct = 0.25, only.pos = TRUE). Statistical significance was determined as a Bonferroni-adjusted p-value <0.05 and an average log2 fold change >0.25. Canonical markers from PanglaoDB and CellMarker were referenced to annotate major cell types^51,52^.

Preprocessed snRNA-seq data for the thoracic aorta was downloaded from https://singlecell.broadinstitute.org/single_cell/study/SCP1265, and cell clusters and annotations from the original study were applied^11^.

To compare cell type expression similarity between arterioles and the thoracic aorta, pseudo-bulk RNA-seq profiles were generated for each cell type within each dataset. DEGs across cell types were calculated using the FindAllMarkers function, with significance determined by a Bonferroni-adjusted p-value < 0.05. Common DEGs across datasets were then used for pairwise Spearman correlation analysis to compare cell types between the two datasets based on their pseudo-bulk expression profiles.

### MAGMA cell typing

Preprocessed study-level GWAS summary statistics for blood pressure regulation and hypertension end-organ damage traits were downloaded from the GWAS Catalog (https://www.ebi.ac.uk/gwas/). This included systolic and diastolic blood pressure (BP) from studies by Keaton JM et al.^53^, Surendran P et al.^54^, and Wojcik GL et al.^55^; pulse pressure from Keaton JM et al.^53^ and Surendran P et al.^54^; essential hypertension from Wojcik GL et al.^55^; stroke from Malik R et al.^56^ and Mishra A et al.^57^; peripheral arterial disease from Sakaue S et al.^58^; diabetic nephropathy from Sakaue S et al.^58^ and Zhou W et al.^59^; and diabetic retinopathy from Backman JD et al.^60^ and Zhou W et al.^59^. Cell type-trait association analysis was performed using the celltype_associations_pipeline from the MAGMA.Celltyping R package (v2.0.11)^61^. The top 10% mode was used to focus on the most enriched genes, as it also has a stronger ability to identify associations and aligns more closely with other gene-trait association methods^62^. Significance was determined using a false discovery rate (FDR) of < 0.05.

### Integrated data analysis

GWAS summary statistics for arteriole-related traits were downloaded from the GWAS Catalog (https://www.ebi.ac.uk/gwas/), including systolic blood pressure (BP), diastolic BP, pulse pressure, essential hypertension, stroke, peripheral arterial disease, diabetic nephropathy, and diabetic retinopathy. Only SNPs with a p-value < 5e-8 were retained for further analysis. To assess whether the percentage of SNPs covered by Micro-C or pan-promoter Capture Micro-C in arterioles and EDAs was significantly higher in arteriolar traits, we selected traits not directly or primarily related to the arteriole and calculated their SNP coverage percentages. These traits included birth weight, bone density, breast cancer, colorectal cancer, Alzheimer’s disease, Parkinson’s disease, multiple sclerosis, rheumatoid arthritis, asthma, psoriasis, and chronic obstructive pulmonary disease. A one-sided binomial test was applied to compare SNP coverage for each arteriolar trait against the average percentage across non-arteriole-relevant traits.

The ClusterProfiler R package (v4.6.2) was used for pathway enrichment analysis on genes associated with arteriole-related SNPs through chromatin interactions captured by pan-promoter Capture Micro-C^63^. Low-abundance genes, defined as those with an average FPKM value less than 0.1, were excluded from the analysis.

Following the same pipeline as before, SNP regulatory annotation was performed using Ensembl annotation^14^. The coverage of SNPs in each regulatory category by promoter Capture Micro-C was calculated separately for promoter bins and distal bins. A Fisher’s exact test was applied to assess whether SNPs were enriched in specific regulatory categories based on their bin location.

To test the enrichment of chromatin-interacting genes in cell type-specific genes, a Fisher’s exact test was applied. Interacting genes were grouped by: 1) interaction types, including promoter-enhancer SNP interacting genes, promoter-interacting genes, and all interacting genes, and 2) the occurrence of interactions by sample, categorized as arteriole-specific, EDA-specific, or overlapping.

Visualization was performed using ggplot2^64^.

### SNP editing in hiPSCs

The generation of isogenic hiPSCs with homozygous low-BP and high-BP alleles for the SNP rs1882961 was achieved using an efficient two-step approach as we described^24^. Briefly, two synthetic sgRNAs targeting genomic regions flanking rs1882961 (**Supplementary Table S15**) and spCas9-2NLS nuclease protein were used to delete a segment of DNA around rs1882961. After 48 hrs of transfection, cells underwent clonal expansion. hiPSC clones showing the intended deletion based on quick extraction of DNA and PCR (**Supplementary Table S15**) were further characterized for the deletion by PCR and sanger sequencing. hiPSC line with the deletion of the region around rs1882961 was tested for their genetic stability by molecular karyotyping using hPSC Genetic Analysis Kit (STEMCELL). For the second step of 2-step SNP editing, homology dependent knock-in approach was used with a gRNA targeting the junction of rs1882961 region deletion site and ssDNA donor fragment for low-BP rs1882961-C or high-BP rs1882961-T alleles with 100-150 bp long homology arms at the 3’ and 5’ sides of rs1882961 (**Supplementary Table S15**). Isogenic hiPSC lines with the reconstitution of homozygous low-BP or high-BP alleles were selected, confirmed, and assessed for their pluripotency, differentiation potential, and genetic stability by molecular karyotyping using hPSC Genetic Analysis Kit and Giemsa-based karyotyping.

### Differentiation of hiPSCs to vascular smooth muscle cells (iVSMCs)

The differentiation was performed as we described^24,65,66^. Briefly, hiPSCs were treated with mTeSR™ plus medium with 10 µM Rock inhibitor Y-27632 (STEMCELL Technologies) for 24 h and then N2B27 medium (Life Technologies) plus 8 µM CHIR99021 (Selleck Chemicals) and 25 ng/ml BMP4 (PeproTech) for 3 days to generate mesoderm cells. For VSMC induction, mesoderm cells were grown with N2B27 medium supplemented with 10 ng/ml PDGF-BB (PeproTech) and 2 ng/ml Activin A (PeproTech) for 2 days, and N2B27 supplemented with 2 ng/ml Activin A and 2 µg/ml Heparin (STEMCELL Technologies) for 5 days. VSMCs were enriched by removing CD144 + cells using CD144 magnetic beads. Enriched VSMC cells were further cultured in N2B27 medium supplemented with 10 ng/ml PDGF-BB, 2 ng/ml Activin A and 1 µM PD032590 (Selleck Chemicals) for 6 days for contractile VSMC formation.

### Real-time PCR

RNA extraction and quantitative real-time PCR analysis were performed as described^67^. Primer sequences are shown in **Supplementary Table S15**. Expression levels of mRNAs were normalized to the endogenous control 18S using ΔΔCt method.

### Region-capture Micro-C and data analysis

The assay and data analysis were performed as described with some modifications^24,68,69^. Briefly, Dovetail® Micro-C kit was used to create Micro-C libraries following manufacturer’s User Guide. Region capture Micro-C libraries were then generated with reagents from Dovetail® targeted enrichment panels. The panel of 80bp probes covers the rs1882961 genomic region and promoters of adjacent protein coding genes. The libraries were sequenced using the Illumina NovaSeq sequencers. Pair-end sequencing reads were cleaned with Trim Galore to remove adaptor and low-quality reads. We used Juicer to generate .hic file and Juicebox for direct visualization^38^.

### Generation of SS-Δrs1882961^−/−^ rat

CRISPR/SpCas9 methods were used to delete a noncoding genomic segment in the Dahl salt-sensitive rat (SS/JrHsdMcwi). The LiftOver tool at UCSC Genome Browser placed the non-coding syntenic region to the human rs1882961 at mRatBN7.2 chr11:15,091,368. SpCas9 (QB3 MacroLab, University of California, Berkely) was mixed with a sgRNA targeting the sequence GATGGCCTGCCAGGACAGAG<u>TGG</u> (protospacer adjacent motif underlined and target base in bold) and injected into fertilized SS/JrHsdMcwi strain rat embryos. A founder was identified harboring a 4-bp deletion (mRatBN7.2 chr11:15,091,367-15,091,370 and was backcrossed to the parental strain to establish a breeding colony. This SS-Δrs1882961^−/−^ strain is registered as SS-Del(11p)3Mcwi (RGDID: 407572511). Heterozygous SS-Δrs1882961^+/−^ breeders were maintained on a purified 0.4% NaCl diet AIN-76A (Dyets, Inc).

### Telemetry measurement of BP

Continuous measurement of BP in conscious, freely moving rats was performed using radiotelemetry as described^70,71^. Briefly, a telemetry transmitter (model HD-S10, Data Systems International) was implanted in the right carotid artery when rats were 7 weeks of age. After one week of recovery, continuous BP monitoring was started. Following three days of stable baseline BP recording, the diet was switched to an AIN-76A diet containing 4% NaCl (high salt diet) for 14 days.

## Supporting information

Suppl figures and list of suppl tables

## AUTHOR CONTRIBUTIONS

YL led omics assays and performed data analysis. RP performed hiPSC and molecular experiments. QQ developed data analysis strategies and analyzed and integrated data. PL analyzed data. HX performed animal experiments. JW collected human arterioles. BT contributed to omics assays. RY contributed to human arteriole collection. KU contributed to animal experiments. MG contributed to the development of the mutant rat. CY contributed to human arteriole collection. MKM contributed to hiPSC experiments. ASG, AWC, SR, and AMG contributed to study design and data interpretation. AMG led the development of the mutant rat. MEW led the collection of human arterioles and contributed to study conception and design. ML conceived and designed the study. QQ, YL, RP, PL, HX, and ML drafted the manuscript. All authors edited and approved the manuscript.

