## Supplementary material for "Chromatin interaction maps of human arterioles reveal new mechanisms for the genetic regulation of blood pressure": Suppl figures and list of suppl tables

[1]Department of Physiology, University of Arizona College of Medicine – Tucson, Tucson, AZ, USA; [2]Molecular Systems Medicine Initiative, University of Arizona Health Sciences, Tucson, AZ, USA; [3]Department of Physiology, Medical College of Wisconsin, Milwaukee, WI, USA; [4]Cardiovascular Center and Department of Medicine, Medical College of Wisconsin, Milwaukee, WI, USA; [5]The Jackson Laboratory, Bar Harbor, ME, USA; [6]Versiti Blood Research Institute, Milwaukee, WI, USA; [7]Department of Pediatrics, Division of Hematology, Oncology, and Transplantation, Medical College of Wisconsin, Milwaukee, WI, USA; [8]Department of Cell Biology, Neurobiology, and Anatomy, Medical College of Wisconsin, Milwaukee, WI, USA.

*co-first authors.

Corresponding author:

Mingyu Liang,

**Supplementary figure legends**

**Supplementary Figure S1. Sizes of chromatin loops identified by Micro-C.** Dash line and number represent the median of the loop size. EDA, endothelium-denuded arteriole.

**Supplementary Figure S2. Distance between interacting chromatin regions based on pan-promoter capture Micro-C analysis.** Dash line and number represent the median distance of the interacting chromatin regions. EDA, endothelium-denuded arteriole.

**Supplementary Figure S3. Comparison of gene expression profiles in intact arterioles, endothelium-denuded arterioles (EDA), and endothelial cells (EC).** N=3.

**Supplementary Figure S4. Expression of marker genes for endothelial cells (A) and vascular smooth muscle cells (B) in tissue components of human arterioles.** EDA, endothelium-denuded arteriole. N=3. Data shown as Mean ± SEM. *, $p < 0.05$, one-way ANOVA followed by Holm-Sidak test.

**Supplementary Figure S5. Comparison of DNA methylation profiles in intact arterioles, endothelia-denuded arterioles (EDA), and endothelial cells (EC).** DMR, differentially methylated regions. N=4.

**Supplementary Figure S6. Correlation between gene expression and methylation in gene promoter regions.** EDA, endothelium-denuded arteriole; EC, endothelial cells.

**Supplementary Figure S7. Most promoter chromatin interacting regions, based on pan-promoter capture Micro-C, contained regulatory elements. A.** Number of chromatin interactions containing regulatory elements in one or both chromatin contact regions. None, one, and both refer to the presence of regulatory elements in none, one, or both of the chromatin contact regions forming a chromatin interaction. **B.** Pan-promoter capture Micro-C analysis indicates that most chromatin regions interacting with gene promoters contain enhancers, other promoters, and transcriptional factor binding sites. Chromatin interactions were categorized as involving promoter-promoter (PP), enhancer-promoter (EP), enhancer-enhancer (EE), enhancer-transcription factor binding site (ET), and promoter-transcription factor binding site (PT), and transcription factor binding site-transcription factor binding site (TT) interactions in human arterioles and EC-denuded arterioles (EDA). The categories were not mutually exclusive. In other words, an interaction was counted in all categories that it fell into.

**Supplementary Figure S8. Generation of isogenic hiPSCs with either homozygous low-BP or high-BP allele of non-coding SNP rs1882961. A.** Two-step genome editing scheme using CRISPR-Cas9. **B.** Sanger sequencing confirmed the deletion of a genomic segment containing rs1882961. Sanger sequencing confirmed the reconstitution of the genomic segment containing either homozygous low-BP (**C**) or high-BP allele (**D**) of rs1882961.

**Supplementary Figure S9. Quality check for reconstituted hiPSC lines with either low-BP or high-BP allele of non-coding SNP rs1882961. A.** Mycoplasma contamination screening for reconstituted hiPSCs. Y1 and Y7, low BP allele cell lines; Z3, Z5, Z16 and Z19, high BP allele cell lines; 39b, original hiPSC line; X85, hiPSC line with deletion of a region containing rs1882961. **B.** Proliferation assay. Equal number of cells were seeded on day 0 in 6cm dish. Cells were collected on day 4 and total cell numbers for reconstituted hiPSCs were compared with original hiPSC cell line. See panel A for cell line designation. **C.** Representative karyograms for low BP (Y7) and high BP (Z3) clones. Chromosomes of 20 proliferating cells were counted. Two to three cells were karyotyped. These cells had a modal number of 46 chromosomes. The sex chromosome constitution indicates normal female. No consistent abnormalities were observed in the chromosomal number or banding patterns.

**Supplementary Figure S10. Comparative mapping and confirmation of deletion in rat. A.** The chromosome 21 region local to rs1882961 is syntenic with a region on rat chromosome 11. **B.** A single guide RNA targeting this sequence was delivered along with SpCas9 to SS rat embryos. **C.** A founder harboring a 4-bp deletion at the target, overlapping the rs1882961 orthologous region was identified and bred to establish a breeding colony. **D.** Sanger sequence confirmation of the 4-bp deletion.

**Supplementary tables (submitted as an Excel file)**

Supplementary Table S1: Information for subjects and samples used for Micro-C, Pan-promoter capture Micro-C, poly(A)-dependent RNA-seq, RRBS, and snRNA-seq.

Supplementary Table S2: Quality control reports for sequencing libraries.

Supplementary Table S3: Micro-C total counts of loops detected by mustach.

Supplementary Table S4: Micro-C loops for arteriole at 4kb resolution.

Supplementary Table S5: Micro-C loops for arteriole at 8kb resolution.

Supplementary Table S6: Micro-C loops for arteriole at 16kb resolution.

Supplementary Table S7: Micro-C loops for endothelium-denuded arteriole (EDA) at 4kb resolution.

Supplementary Table S8: Micro-C loops for endothelium-denuded arteriole (EDA) at 8kb resolution.

Supplementary Table S9: Micro-C loops for endothelium-denuded arteriole (EDA) at 16kb resolution.

Supplementary Table S10: Pan-promoter capture Micro-C total counts of chromatin interactions detected by CHiCAGO.

Supplementary Table S11: Pan-promoter capture Micro-C chromatin interactions for arteriole at 10kb resolution.

Supplementary Table S12: Pan-promoter capture Micro-C chromatin interactions for arteriole at 20kb resolution.

Supplementary Table S13: Pan-promoter capture Micro-C chromatin interactions for endothelium-denuded arteriole (EDA) at 10kb resolution

Supplementary Table S14: Pan-promoter capture Micro-C chromatin interactions for endothelium-denuded arteriole (EDA) at 20kb resolution

Supplementary Table S15: Sequences of gRNAs, ssODN, and PCR primers.

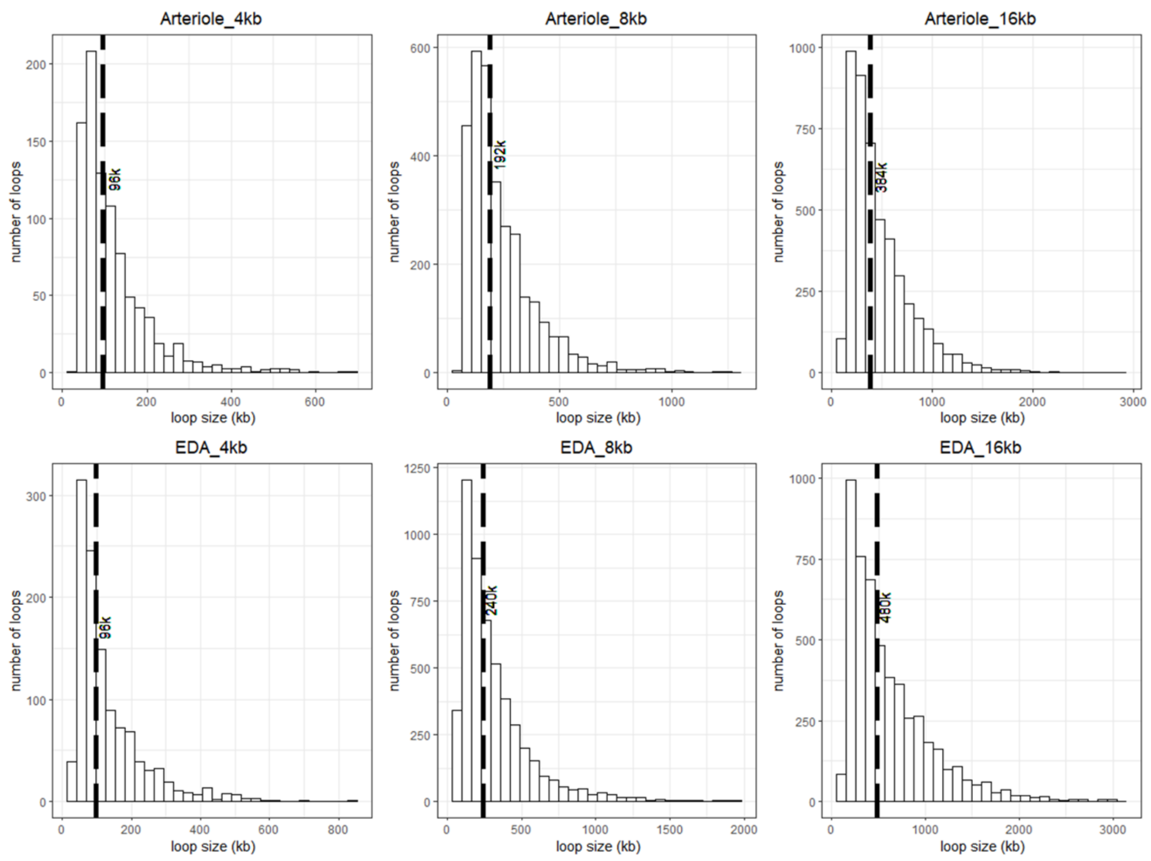


**Supplementary Figure S1. Sizes of chromatin loops identified by Micro-C.** Dash line and number represent the median of the loop size. EDA, endothelium-denuded arteriole.

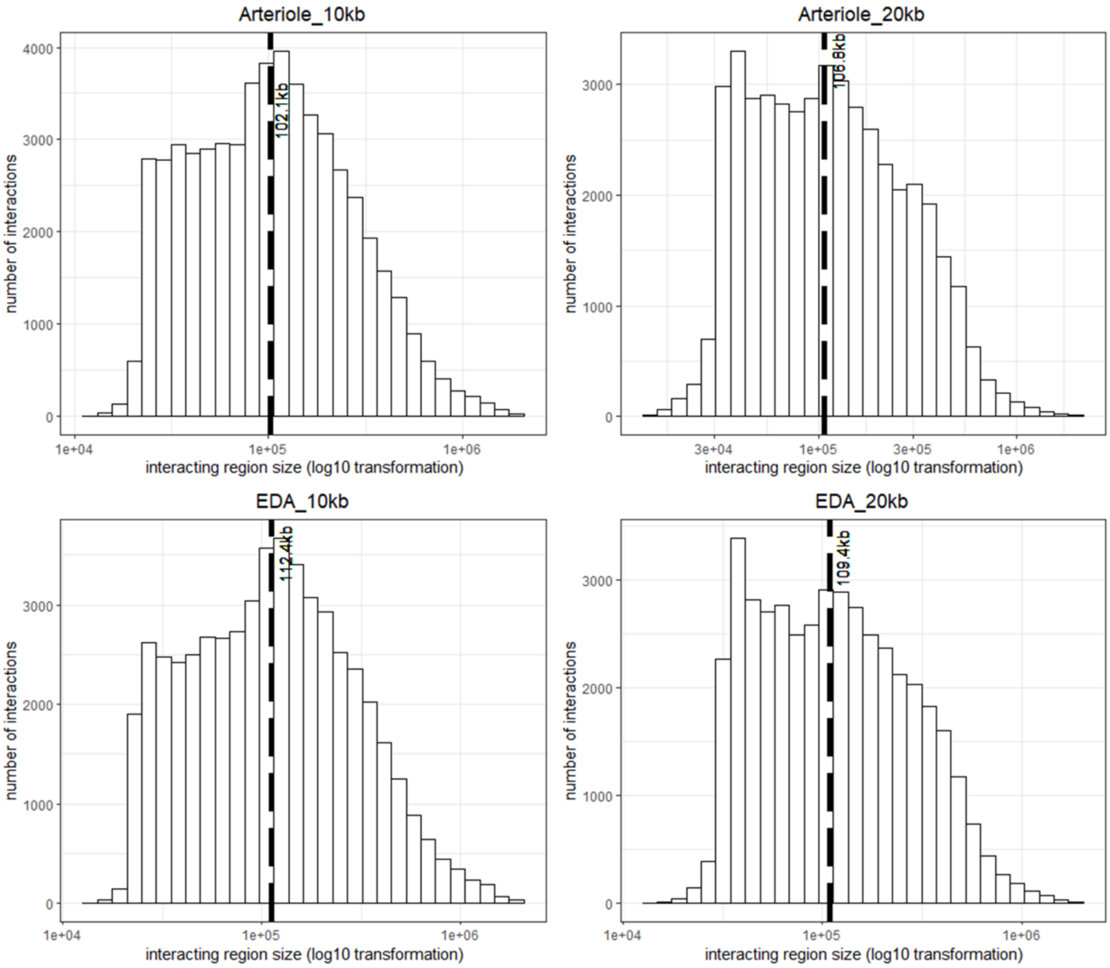


**Supplementary Figure S2. Distance between interacting chromatin regions based on pan-promoter capture Micro-C analysis.** Dash line and number represent the median distance of the interacting chromatin regions. EDA, endothelium-denuded arteriole.

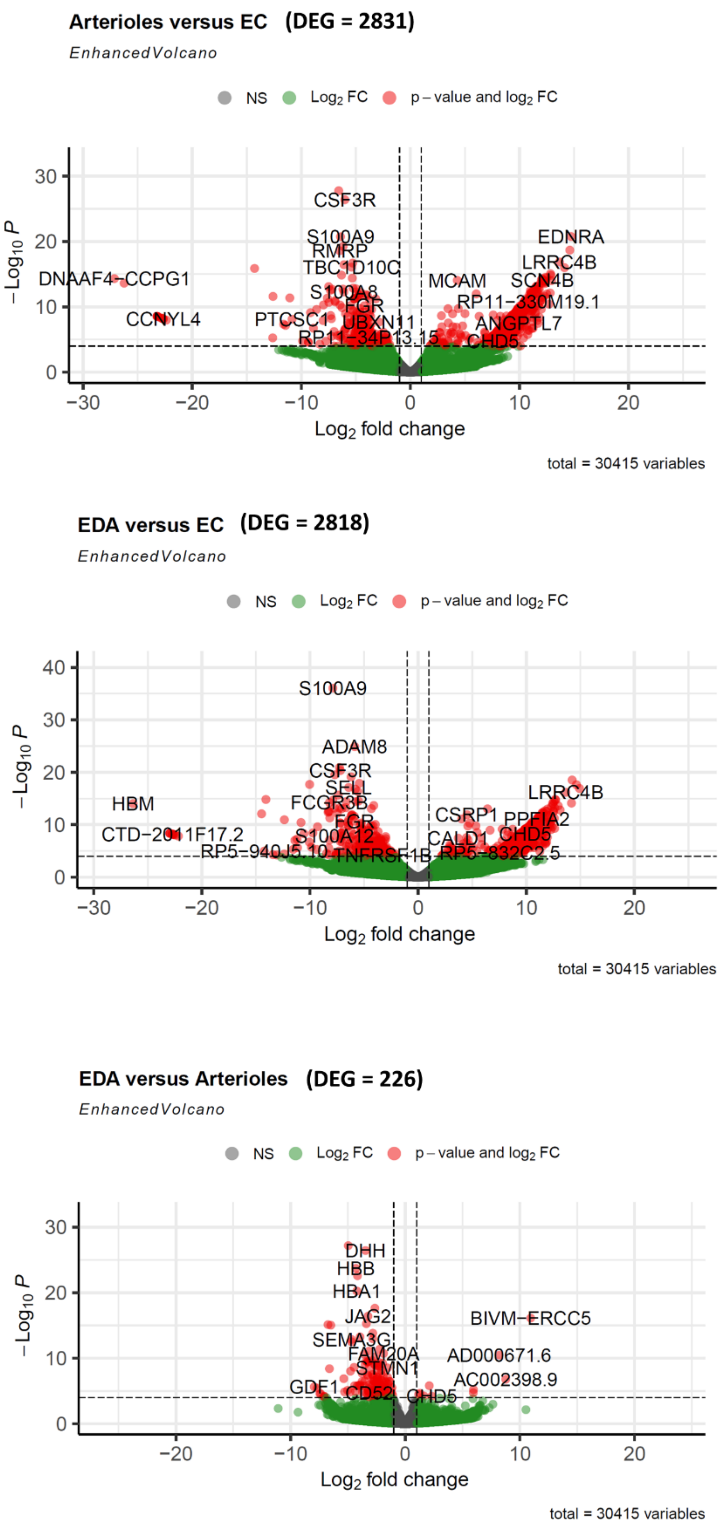


**Supplementary Figure S3. Comparison of gene expression profiles in intact arterioles, endothelium-denuded arterioles (EDA), and endothelial cells (EC).** N=3.

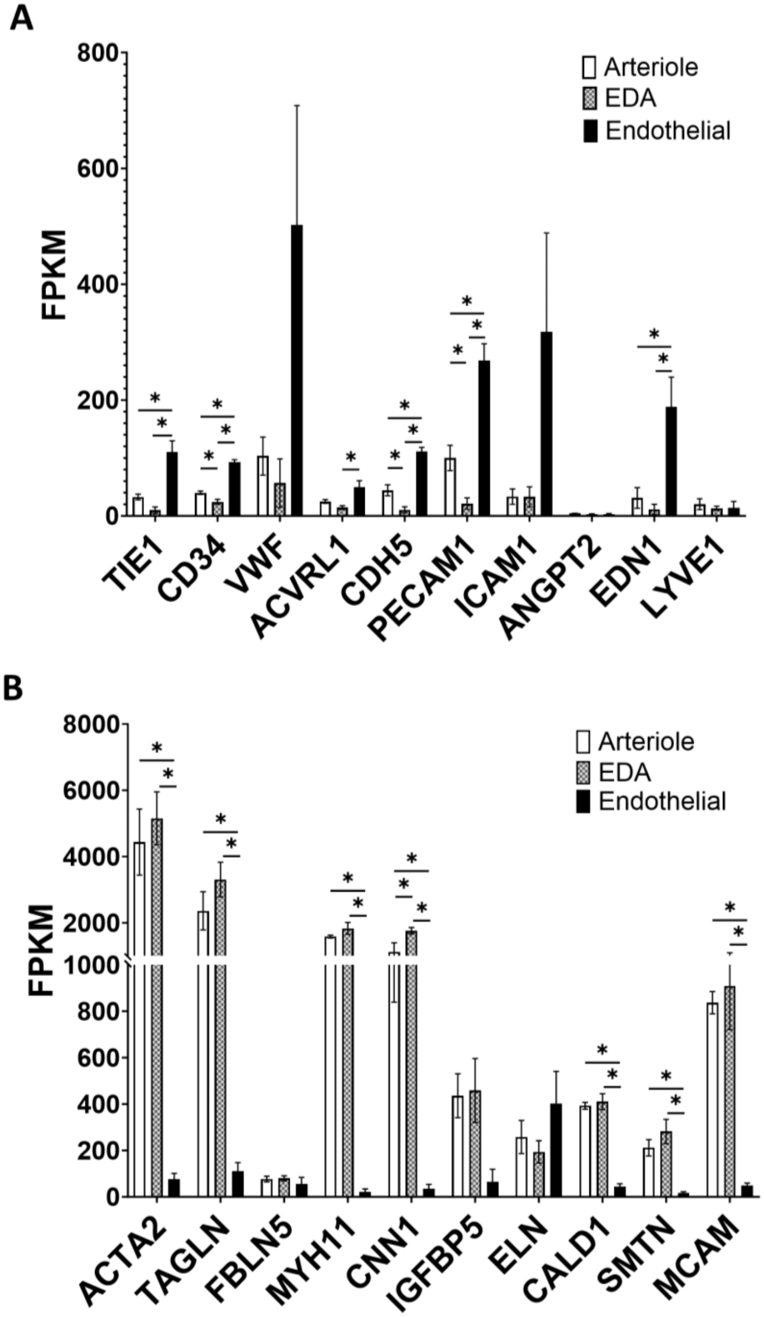


**Supplementary Figure S4. Expression of marker genes for endothelial cells (A) and vascular smooth muscle cells (B) in tissue components of human arterioles.** EDA, endothelium-denuded arteriole. N=3. Data shown as Mean ± SEM. *, $p < 0.05$, one-way ANOVA followed by Holm-Sidak test.

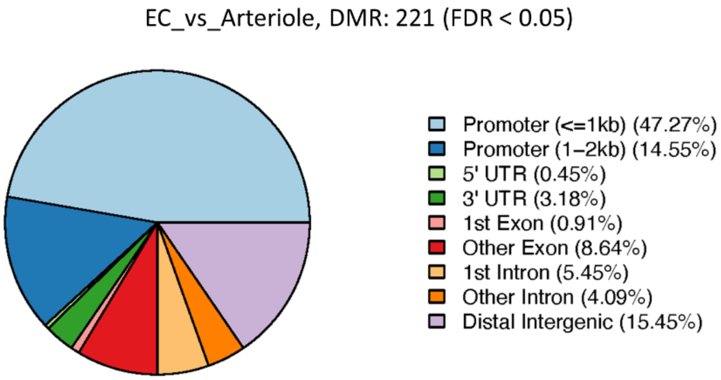


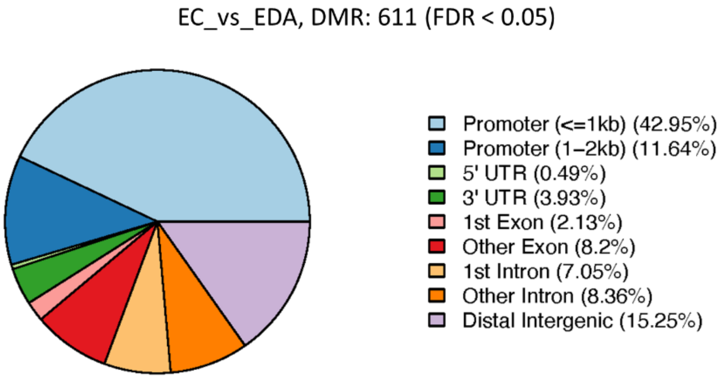


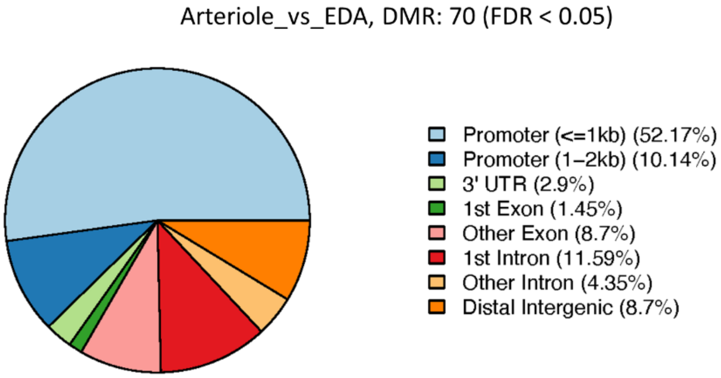


**Supplementary Figure S5. Comparison of DNA methylation profiles in intact arterioles, endothelia-denuded arterioles (EDA), and endothelial cells (EC).** DMR, differentially methylated regions. N=4.

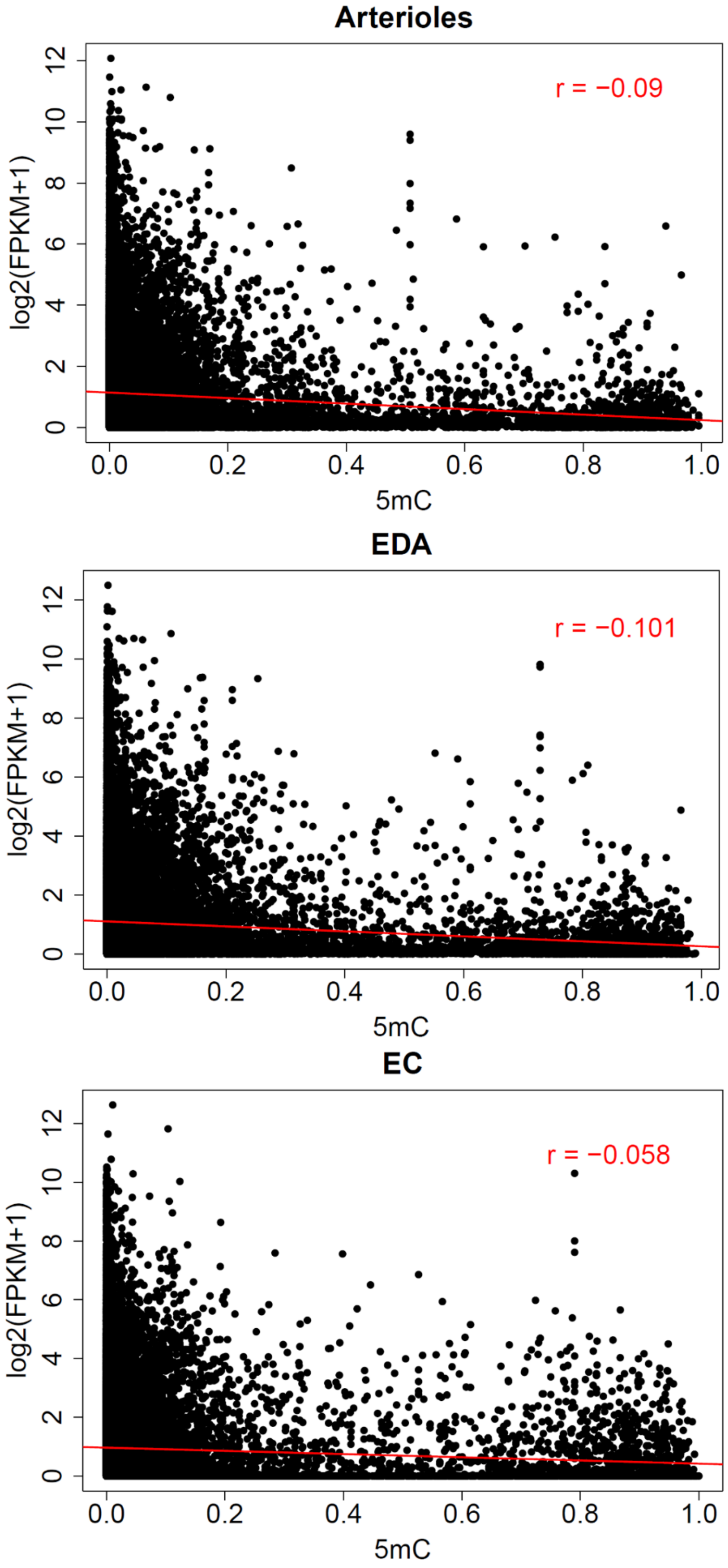

Arterioles
r = −0.09
log2(FPKM+1)
5mC
EDA
r = −0.101
log2(FPKM+1)
5mC
EC
r = −0.058
log2(FPKM+1)
5mC

**Supplementary Figure S6. Correlation between gene expression and methylation in gene promoter regions.** EDA, endothelium-denuded arteriole; EC, endothelial cells.

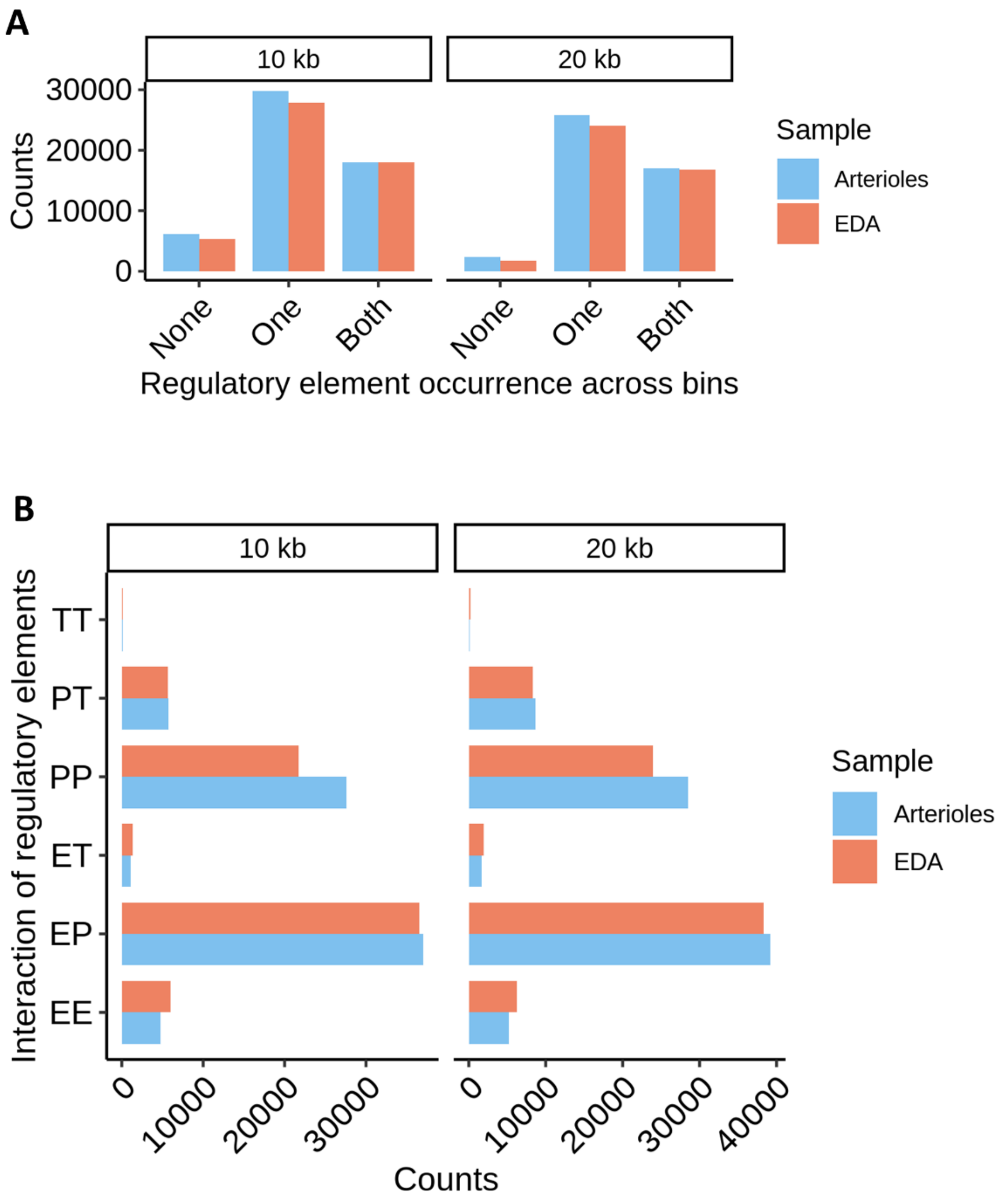


**Supplementary Figure S7. Most promoter chromatin interacting regions, based on pan-promoter capture Micro-C, contained regulatory elements. A.** Number of chromatin interactions containing regulatory elements in one or both chromatin contact regions. None, one, and both refer to the presence of regulatory elements in none, one, or both of the chromatin contact regions forming a chromatin interaction. **B.** Pan-promoter capture Micro-C analysis indicates that most chromatin regions interacting with gene promoters contain enhancers, other promoters, and transcriptional factor binding sites. Chromatin interactions were categorized as involving promoter-promoter (PP), enhancer-promoter (EP), enhancer-enhancer (EE), enhancer-transcription factor binding site (ET), and promoter-transcription factor binding site (PT), and transcription factor binding site-transcription factor binding site (TT) interactions in human arterioles and EC-denuded arterioles (EDA). The categories were not mutually exclusive. In other words, an interaction was counted in all categories that it fell into.

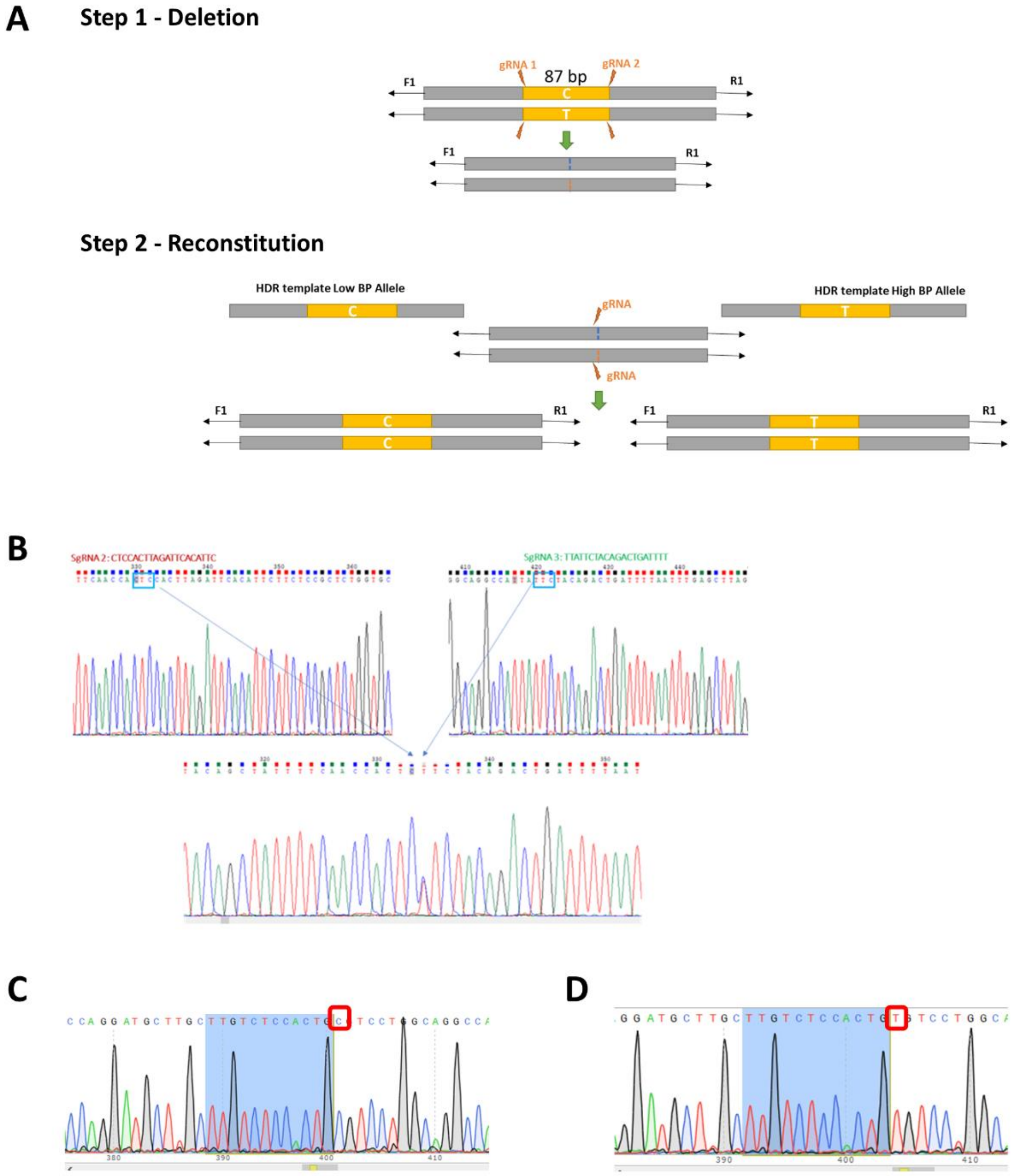


**Supplementary Figure S8. Generation of isogenic hiPSCs with either homozygous low-BP or high-BP allele of non-coding SNP rs1882961. A.** Two-step genome editing scheme using CRISPR-Cas9. **B.** Sanger sequencing confirmed the deletion of a genomic segment containing rs1882961. Sanger sequencing confirmed the reconstitution of the genomic segment containing either homozygous low-BP (**C**) or high-BP allele (**D**) of rs1882961.

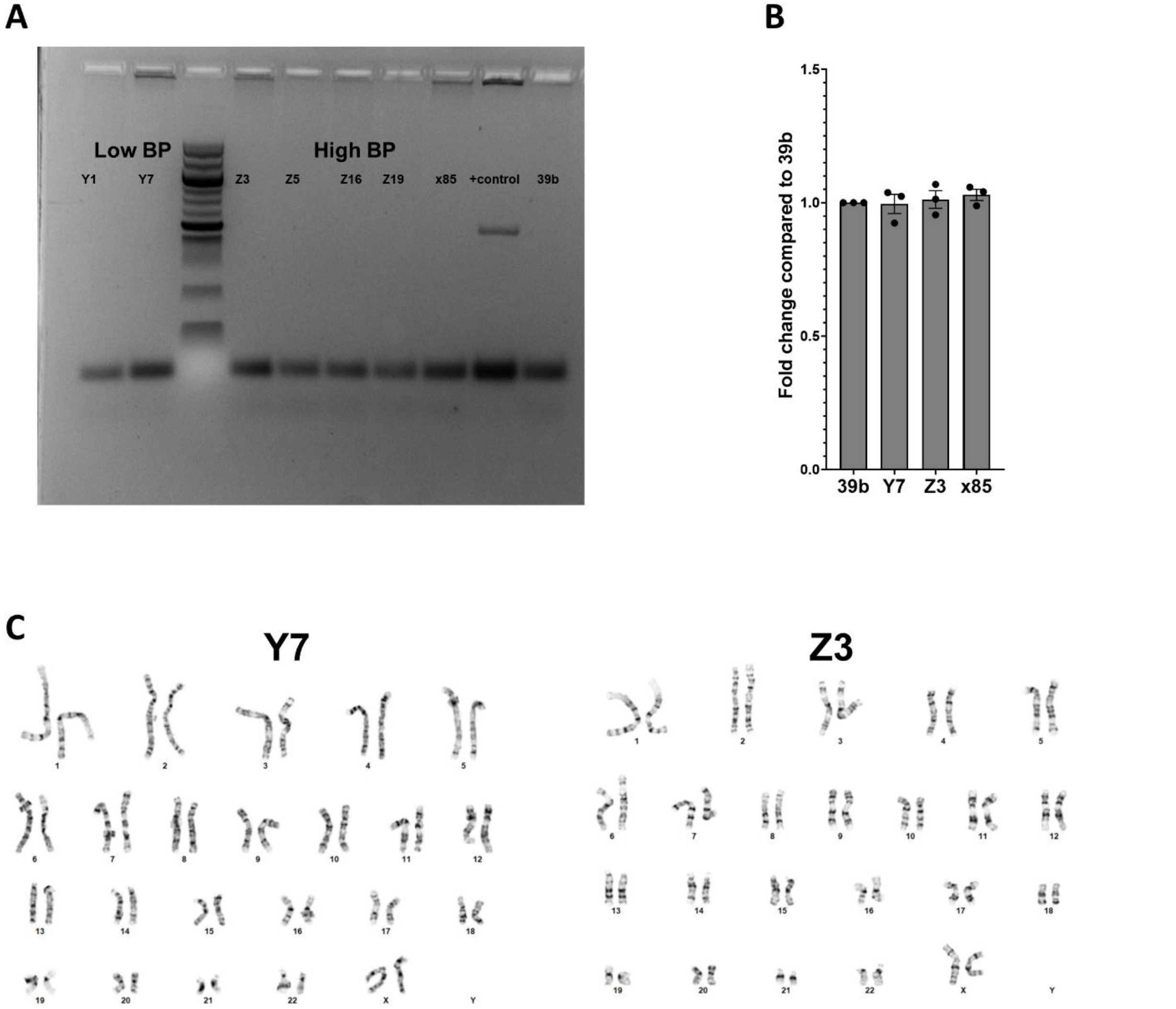


**Supplementary Figure S9. Quality check for reconstituted hiPSC lines with either low-BP or high-BP allele of non-coding SNP rs1882961. A.** Mycoplasma contamination screening for reconstituted hiPSCs. Y1 and Y7, low BP allele cell lines; Z3, Z5, Z16 and Z19, high BP allele cell lines; 39b, original hiPSC line; X85, hiPSC line with deletion of a region containing rs1882961. **B.** Proliferation assay. Equal number of cells were seeded on day 0 in 6cm dish. Cells were collected on day 4 and total cell numbers for reconstituted hiPSCs were compared with original hiPSC cell line. See panel A for cell line designation. **C.** Representative karyograms for low BP (Y7) and high BP (Z3) clones. Chromosomes of 20 proliferating cells were counted. Two to three cells were karyotyped. These cells had a modal number of 46 chromosomes. The sex chromosome constitution indicates normal female. No consistent abnormalities were observed in the chromosomal number or banding patterns.

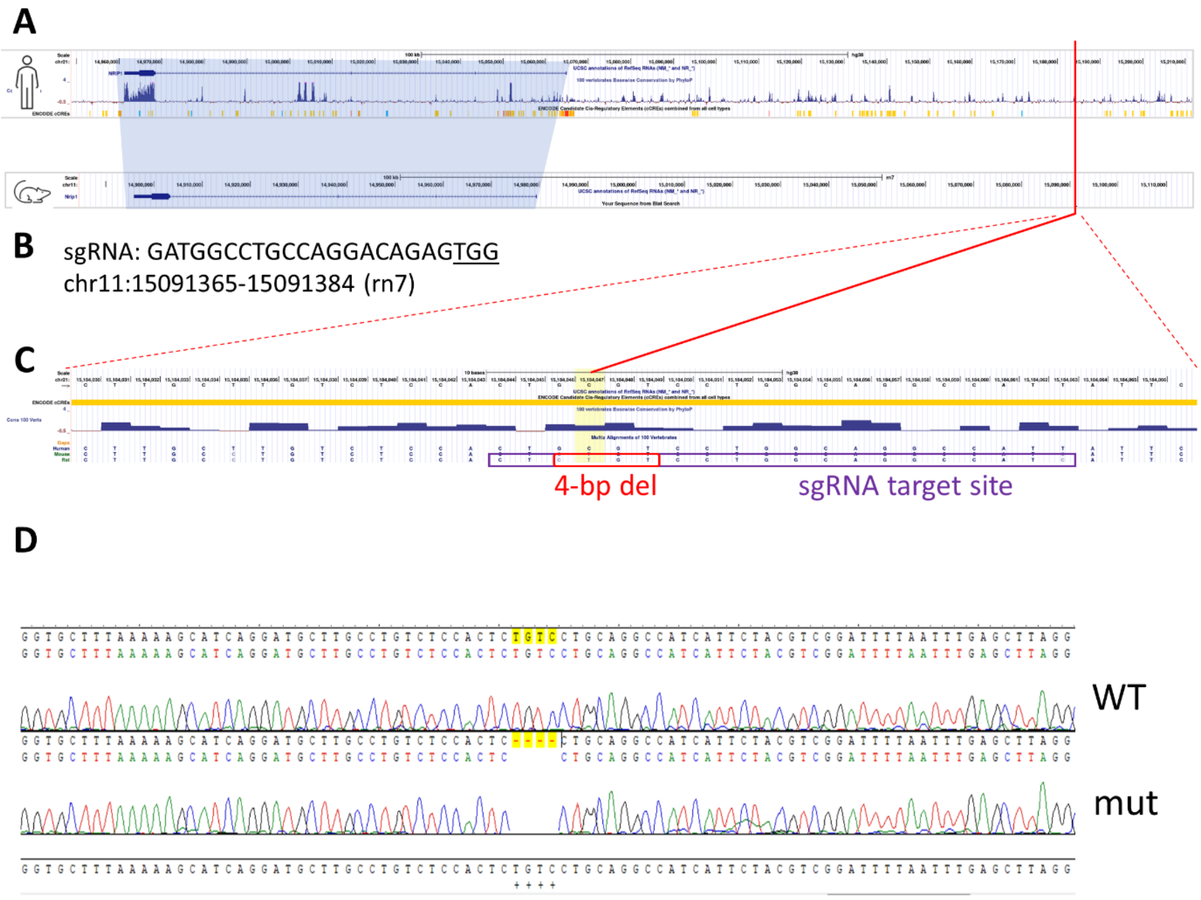


**Supplementary Figure S10. Comparative mapping and confirmation of deletion in rat. A.** The chromosome 21 region local to rs1882961 is syntenic with a region on rat chromosome 11. **B.** A single guide RNA targeting this sequence was delivered along with SpCas9 to SS rat embryos. **C.** A founder harboring a 4-bp deletion at the target, overlapping the rs1882961 orthologous region was identified and bred to establish a breeding colony. **D.** Sanger sequence confirmation of the 4-bp deletion.
